# Early precursor-derived pituitary gland tissue-resident macrophages play a pivotal role in modulating hormonal balance

**DOI:** 10.1101/2024.05.07.592305

**Authors:** Henna Lehtonen, Heli Jokela, Julian Hofmann, Lauriina Tola, Arfa Mehmood, Florent Ginhoux, Burkhard Becher, Melanie Greter, Marko Salmi, Heidi Gerke, Pia Rantakari

## Abstract

The pituitary gland is the central endocrine regulatory organ underneath the brain, producing and releasing a variety of hormones that coordinate major body functions. The physical location of the pituitary gland underneath the brain, though outside the protective blood-brain barrier, leads to a unique immune environment of the pituitary that has not been studied. Here, we defined the development, diversity, spatial niche, and origin of the pituitary gland macrophage subsets using single cell transcriptomics, fate mapping, and imaging. We identified early yolk sac precursors solely seeding pituitary gland macrophages which are maintained by proliferation. Macrophage depletion experiments unveiled the essential contribution of early macrophages in the pituitary gland’s hormonal production and in modulating the post-pubertal expression of genes related to the sexually dimorphic processes regulated by the pituitary gland. Altogether, these findings provide novel information on pituitary gland macrophages and advance our understanding of immune-endocrine system crosstalk.

## Introduction

The pituitary gland is a central endocrine regulatory organ located underneath the brain. Being connected to the hypothalamus via a stalk including blood vessels and nerves, the pituitary gland acts as a signal mediator between the hypothalamus and peripheral organs. The function of the pituitary gland is to produce and release hormones and, thus, coordinate several central physiological pathways that impact postnatal growth, puberty, lactation, reproduction, stress response, water balance, and immune response ^1–3^.

The pituitary gland comprises two ectodermal structures ^1,4^. The neural ectoderm gives rise to the posterior lobe (neurohypophysis), responsible for storing and secreting arginine vasopressin (AVP) and oxytocin (OXT), produced by the hypothalamus. The posterior lobe consists of the stalk and the neural lobe and is composed of axons and pituicytes, glial cells surrounding and supporting hormonal secretion from these axonal endings ^5^. The surface ectoderm, which creates Rathke’s pouch, is the precursor to the anterior and intermediate lobes (adenohypophysis and intermedia, respectively). The intermediate lobe is formed of melanotrophs that produce the melanocyte-stimulating hormone, whereas the anterior lobe is composed of five specialized endocrine cell types producing and secreting different hormones and non-endocrine cells such as follicular-stellate (FS) cells, supporting and coordinating endocrine cell function, vascular endothelial and immune cells ^3,4,6^.

Macrophages are critical regulators of tissue homeostasis ^7–9^. Tissue-resident macrophages generally originate from embryonic precursors (yolk sac and/or fetal liver) and are maintained locally through self-renewal ^10,11^. Under homeostatic conditions, each adult organ has its own ontogenetically and functionally distinct macrophage pool controlling tissue-specific and niche-specific functions ^10,12–18^. Particularly noteworthy among macrophages are central nervous system macrophages, microglia, and border-associated macrophages (BAMs). These cells are predominantly of embryonic origins and sustain themselves through proliferation ^19^, and only during the aging process and during neuronal diseases can BAMs be renewed through monocyte recruitment^20,21^. Diverse roles played by macrophages have also been described in endocrine organs ^22–28^. Macrophages in pancreatic islets are vital players in metabolic inflammation, underlining the development of insulin resistance ^22^. Whereas in the ovary, they play an essential role in ovarian folliculogenesis and ovulation ^23,29^, and in the male reproductive tract, they regulate vascularization and spermatogenesis ^24,30,31^.

In the pituitary gland, macrophages form the foremost immune cell population ^32,33^. However, the origin, features, or their relationship to pituitary gland function in homeostasis or pathological conditions are not understood. Herein, we constructed a high-density transcriptional map elucidating the intricacies of pituitary gland macrophages (pitMØ) and unveiled their developmental kinetics and distinctive characteristics. We unveiled a postnatal divergence within the initially homogeneous pitMØ population. Notably, these subsets exhibited marked distinctions in their gene expression profiles and spatial localization, shedding light on the nuanced dynamics shaping the development of pituitary gland macrophages. Within pitMØ, we identified microglia-like macrophages in the posterior lobe and vascular-associated macrophages in the anterior lobe of the pituitary gland. Furthermore, we revealed that the pitMØ originate from the early yolk sac precursors and relies on proliferation for self-maintenance. Macrophage depletion unveiled a hitherto unknown contribution of macrophages to the homeostatic development of the pituitary gland, as depletion altered the post-pubertal expression of genes related to the sexually dimorphic processes and calcium signaling and led to abnormal pituitary hormonal levels. Altogether, our data implicate a novel role for macrophages in regulating pituitary gland development and its core functions.

## Results

### At a steady state, the pituitary gland hosts transcriptionally distinct macrophage subsets

To gain a comprehensive view of macrophage subsets that reside within the developing homeostatic pituitary gland, we collected the whole pituitary glands from newborn (NB) to adult (8 weeks of age) wild-type (WT) mice, isolated and enriched total CD45^+^ leukocytes by fluorescent-activated cell sorting (FACS) to utilize a droplet-based single cell transcriptomics approach (10× Chromium; Figure 1A). ScRNA sequencing (scRNA-seq) analyses revealed the complex immune environment in the developing pituitary gland, including macrophages, monocytes, T and B cells, natural killer (NK) cells, neutrophils, dendritic cells (DCs) and mast cells (Figures S1A-S1C). We then performed additional subclustering of macrophage populations expressing canonical macrophage markers (*Fcgr1*, *Adgre1*, and *Cd68*) and monocyte populations expressing *Itgam* and either *Ly6c2* or *Nr4a1*. Low-quality clusters and clusters containing mixed cell types were excluded from this analysis (Figures S1B and S1D). The subclustering approach allowed for a more detailed assessment of the emergence and phenotypic profile of these specific cell subsets within the developing pituitary gland. (Figures S1C and S1E). Subclustering of macrophage and monocyte cells led to the identification of six macrophages and two monocyte clusters (Figure 1B). Cluster frequencies presented a kinetical progression and fluctuation over time, and there was a dramatic increment in the total pitMØ numbers after birth (Figures 1C, D, and S1F).

**Figure 1.**
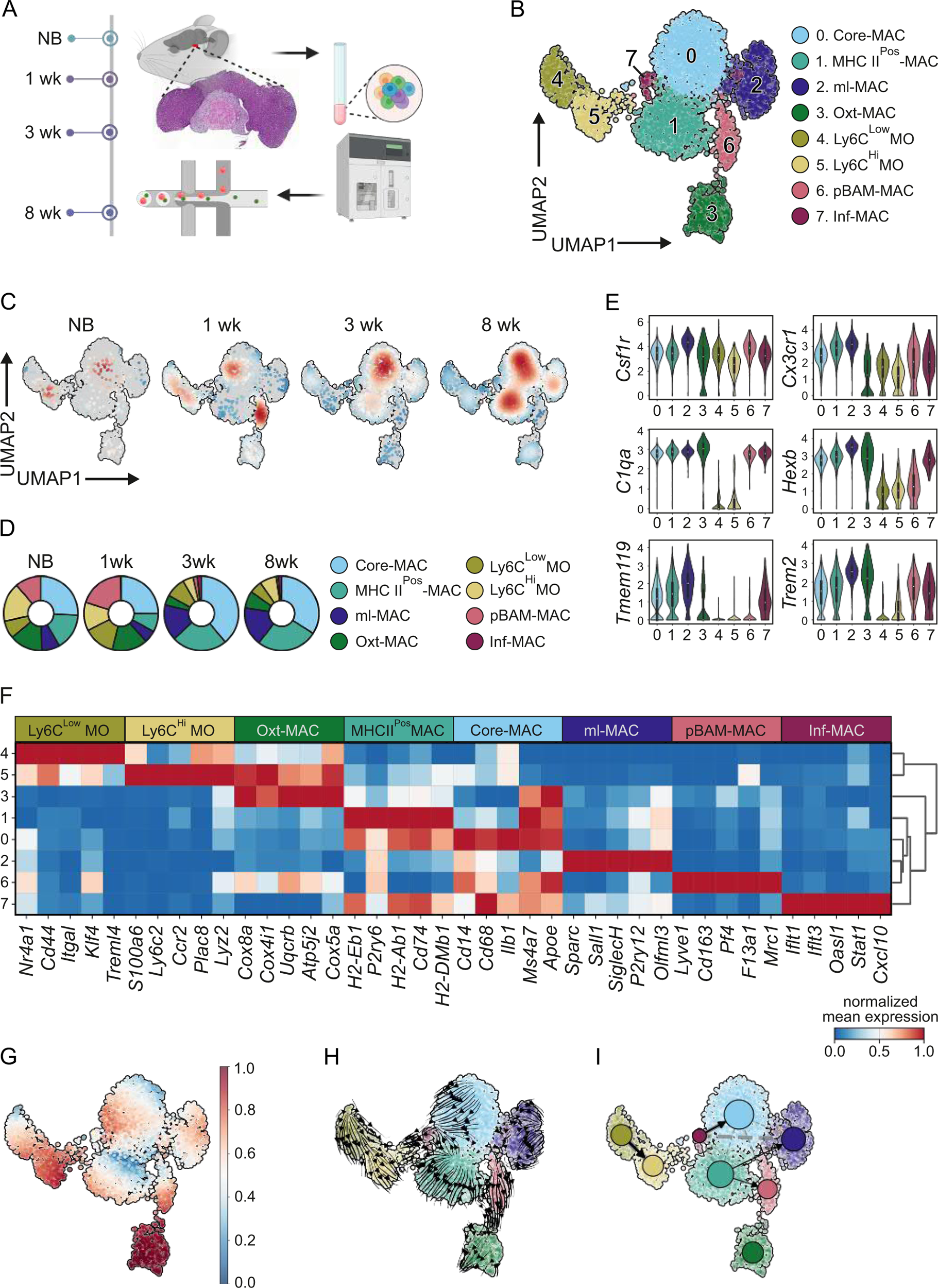
At a steady state, the pituitary gland hosts macrophage and monocyte subsets that exhibit distinct transcriptional variability. (A) Workflow. Whole pituitary glands from newborn (NB), 1-, 3-, and 8-week-old mice were used for this study. Pituitary glands were digested, and CD45^Pos^ cells were enriched by fluorescence-activated cell sorting. 10X Genomics Chromium scRNA-seq was used to sequence the cells. Created with BioRender.com. (B) Uniform manifold approximation and projection (UMAP) plot of mouse pituitary gland macrophages (pitMØ; cells expressing *Fcgr1*, *Adgre1*, and *Cd68*) and monocytes (cells expressing *Ly6c2*) colored by the cluster. (C) UMAP plots highlighting the cell embedding across timepoints. Color intensity represents cell density from low (blue) to high (red). (D) Donut charts present the cell frequency of pitMØ and monocytes in the developing pituitary gland. (E) Violin plots showing expression of selected genes in pitMØ and monocyte clusters. (F) Heatmap of single-cell expression of selected differentially expressed genes in pitMØ and monocyte clusters. The color code indicates the expression level from low (blue) to high (red). (G) CytoTRACE-predicted transcriptional dynamics of pitMØ and monocytes on the UMAP. The color code indicates the computed cell differentiation level from less (blue) to highly differentiated (red). (H) Differentiation velocity field of pitMØ and monocytes on the UMAP. Arrow lines indicate putative developmental differentiation trajectories. (I) Partition-based graph abstraction (PAGA) analysis with pseudotime-directed edges of pitMØ and monocytes projected on the UMAP with node connectivities (dashed) and directed transitions (solid/arrows).

All macrophage clusters were uniformly expressing *Csf1r*, *Cx3cr1,* and *C1qa,* and the majority were also highly expressing *Hexb, Tmem119,* and *Trem2*, typical markers for microglia (Figure 1E). Macrophage sub-clusters 0 and 1 were predominant in all stages and exhibited similar overlapping transcriptomic profiles (Figures 1D, 1F, S1F, and S1G). Cluster 0 (hereafter called Core-MAC) mainly expressed common macrophage markers (e.g., *Cd14, Cd86,* and *Il1b*; Figure 1F and Table S1), whereas cluster 1 (hereafter called MHC II^Pos^-MAC) showed increased expression of immune response activation and antigen presentation genes such as *H2-Eb1*, *P2ry6*, *H2-Ab1*, *Cd74,* and *H2-DMb1*; (Figure 1F and Table S1). Interstingly, Cluster 2 (hereafter called microglia-like MAC; ml-MAC) was enriched in microglia signature genes such as *Sparc*, *Sall1*, *Siglech*, *P2ry12*, *Olfml3*, *Fscn1,* and *Nav3* ^34–37^ and expressed higher levels of *Mertk* than any other macrophages in the pituitary gland (Figure 1F and Table S1). On the other hand, cluster 6 was expressing a prototypical BAM (hereafter called pBAM-MAC) signature gene such as *Lyve1*, *Cd163*, *Pf4*, *F13a1, Fcna*, *Folr2*, *Dab2, Cbr2*, *Mrc1*, *Clec10a,* and *Stab1* ^13,37,38^ (Figures 1F and Table S1). From these genes, *Lyve1*, *Mrc1*, and *F13a1* have previously been reported to be markers of vascular-associated tissue-resident macrophages ^14,39^. The abundance of pBAM-MAC cells peaked at 1 week of age, from where it diminished towards adulthood almost entirely (Figures 1C, D, and S1F). Cluster 3 (hereafter called Oxt-MAC) had a relatively high enrichment of genes related to oxidative phosphorylation (e.g., *Cox8a*, *Cox4i1, Uqcrb, Atp5j2*, *Cox5a,* and *Cox6b1*) (Figure 1F and Table S1) and showed increased expression of genes involved in ribosomal biogenesis and protein synthesis in activated macrophages (Table S1). The small cluster 7 (Inf-MAC) only appeared after one week of age and expressed genes related to response to interferon beta and monocyte differentiation (e.g., *Ifit1*, *Ifit3, Oasl1, Stat1, Mndal*, *Gbp5,* and *Cxcl10*; Figures 1C, F and Table S1). However, the proportion and total cell number of cluster 7 did not markedly increase between juvenile and adult animals (Figures 1D and S1F). We furthermore noticed that Core-Mac, ml-Mac, pBAM-Mac, and Inf-Mac were also characterized by a high expression of immediate early genes (e.g., *Jun*, *Junb*, *Jund*, *Fos*, *Egr1*, and *Klf6*) encoding transcription factors that may define a subpopulation of transcriptionally active cells (Figure S1H and Table S1). A noteworthy observation was that *H2-Ab1* (MHC II) expression increased in all the other macrophage populations except ml-MAC (Figure S1I) as the animals aged. Clusters 4 and 5 were enriched with typical monocyte markers (Figure 1F). Cluster 5 expressed high levels of *S100a6*, *Ly6c2*, *Ccr2*, *Lyz2,* and *Plac8* and was defined as classical monocytes (Ly6C^Hi^MO). On the contrary, cluster 4 showed the upregulation of *Nr4a1*, *Cd44*, and *Itgal*, markers used to identify non-classical monocytes (Ly6C^Low^MO). Additionally, the Ly6C^Low^MO cluster expressed features such as *Klf4*, a regulator of monocyte differentiation, and *Treml4*, a commitment marker of monocytes differentiating to macrophages ^40^ (Figure 1F and Table S1).

To examine potential developmental trajectories of the monocyte/macrophage subtypes, we next applied CytoTRACE ^41^ to predict cell lineage differentiation states of clusters based on the fact that the number of genes expressed in a cell decreases as the cell becomes more differentiated. Pseudotime differentiation trajectory analysis showed that Oxt-MAC cells were the most differentiated, while Core-MACs and the MHC II^Pos^-MACs were the less differentiated clusters. Interestingly, pBAM-MAC cells were in intermediate transitions between the differentiation stages (Figure 1G). We next visualized the directed transition matrix on a UMAP representation with original macrophage/monocyte cluster annotations to recapitulate the central trajectory trends (Figure 1H). This analysis further suggested that Core-MACs and MHC II^Pos^-MACs likely serve as origin clusters to pBAM-MACs, which would eventually differentiate towards Oxt-MAC cells (Figure 1H). Likewise, vectors in Oxt-MAC in the transition matrix pointed away from other clusters, supporting its terminal state status (Figure 1H). Interestingly, the ml-MACs appeared to be an independent and distinct cellular state. No clear transition was evident from the other macrophage or monocyte subpopulations to the ml-MACs cells, predicting the possibility that ml-MACs can originate from the independent origin or differentiation lineage (Figure 1H). Furthermore, the transition matrix from the ml-MACs did not point toward any other clusters (Figure 1H). Monocyte subpopulations (Ly6C^Hi^MO and Ly6C^Low^MO) emerged as distinct entities outside the central trajectory. Intriguingly, these monocyte populations did not exhibit directional vectors toward the macrophage lineages (Figure 1H).

We further utilized the directed partition-based graph abstraction (PAGA) analysis to validate the connectivity among the macrophage and monocyte clusters ^42^. The PAGA analysis recapitulated the central trajectory trends and indicated connectivity between Core-MAC and MHC II^Pos^-MACs and separated Oxt-MACs (Figure 1I). Notably, in the PAGA analysis, the pBAM-MAC cluster appeared to be in an intermediate state, which connected Core-MAC and MHC II^Pos^-MAC clusters to Oxt-MACs. Interestingly, we also found connectivity between Inf-MAC and ml-MACs that was not evident in the transition matrix analyses (Figure 1H). Ly6C^Hi^MO and Ly6C^Low^MO populations emerged as clear outliers, as they deviated significantly from the central trajectory, which was consistent with transition matrix analyses (Figures 1H and I). Collectively, our data reveal the coexistence of multiple diverse macrophage populations, which dramatically increased during the time course. The gradual changes in gene expression of macrophage clusters could reflect more of the different transition stages than separated macrophage populations. Moreover, data analyses suggest that the monocyte precursors may not considerably contribute to pitMØ development.

### Tissue growth induces modification in the phenotype of pituitary gland macrophages

We next used the transcriptomic information to distinguish macrophages with flow cytometry. Expression of CD11b, Ly6C, and F4/80 allowed the resolution of total macrophages and two monocyte subsets, presenting both tissue-resident and possible blood-circulating monocytes (Figure 2A; see the gating strategy in S2A). The pitMØ cell numbers increased manifold when the growth of the pituitary gland is most significant, between the ages of P14 and 5 weeks. Both monocyte subsets, the Ly6C^Hi^MO (Ly6C^Hi^CX3CR1^Low^; classical monocytes) and Ly6C^Low^MO (Ly6C^Low^CX3CR1^Hi^; non-classical monocytes) stayed at constant low cell amount and frequency levels throughout all time points studied (Figure 2A and S2B). An F4/80^Pos^Ly6C^Neg^ pitMØ population, expressing the common macrophage markers CD64 and CX3CR1, was found throughout the development (Figure S2C).

**Figure 2.**
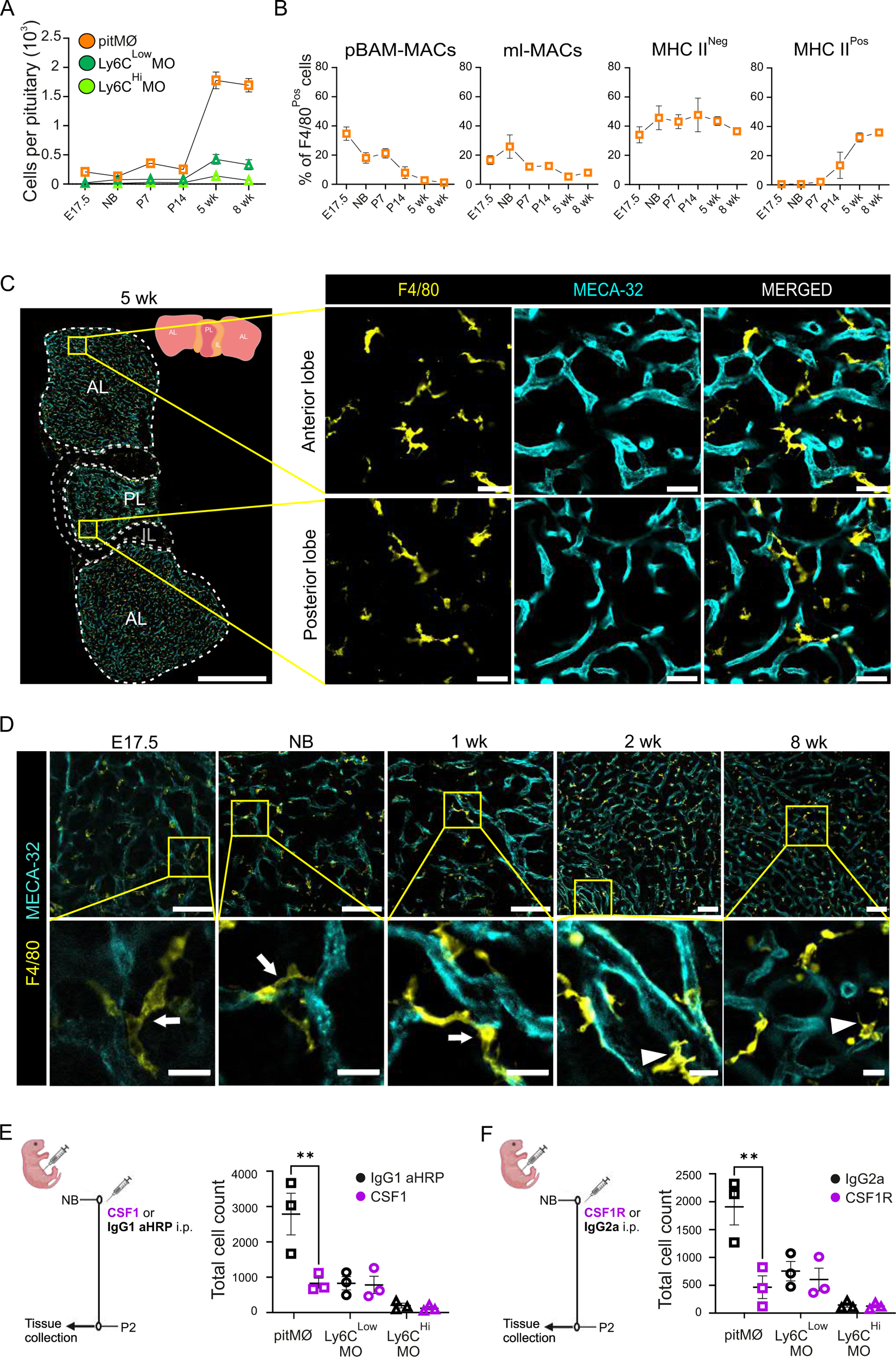
Development modifies the phenotype of CSF1R-dependent pituitary gland macrophages. (A) The cell amount of pitMØ, Ly6C^Low^MO and Ly6C^Hi^MO cells in the WT mouse pituitary gland at indicated timepoints. The quantitative data are shown as mean ± SEM. Each dot represents data from mice pooled together in E17.5 (n=8-12 samples/pool), NB (n=7-11 samples/pool), 1 wk (n=4-6 samples/pool), 2 wk (n=2-5 samples/pool), and from individual mice at 5 wk (n=15 mice) and 8 wk (n=13 mice). Data are from 6-8 individual experiments. (B) Frequency of prototypical BAMs (pBAMs), microglia-like MACs (ml-MACs), MHC II^Neg^, and MHC II^Pos^ cells in F4/80^Pos^ cells in WT pituitary gland at indicated timepoints. The quantitative data are shown as mean ± SEM. Each dot represents data from mice pooled together in E17.5 (n=9-12 samples/pool), NB (n=8-9 samples/pool), 1 wk (n=4-6 samples/pool), 2 wk (n=4-5 samples/pool), and from individual mice at 5 wk (n=8 mice) and 8 wk (n=9 mice). Data are from 3-4 individual experiments. (C) Pituitary gland whole mount immunofluorescence with F4/80 and endothelial marker, MECA-32, from WT pituitary gland of 5-week-old mouse. Inserts are higher magnifications from the boxed anterior lobe and posterior lobe areas. For overview image, scale is 500 µm, and for inserts, 20 µm. Shown are representative images from ≤ 3 independent experiments. (D) Pituitary gland whole mount immunofluorescence with F4/80 and endothelial marker, MECA-32, from WT pituitary glands of E17.5, NB, 1-, 2- and 8-week-old mice. Inserts are higher magnifications from the boxed areas. Arrows indicate a more rounded phenotype at early time points and arrowheads indicate morphology of mature phagocytes at 2-to 8-week-old mice. For overview images, scales are 50 µm, and for inserts, scales are 10 µm. Shown are representative images from ≤ 3 independent experiments. (E) Experimental outline for studying pituitary macrophages and monocytes at postnatal day (P) 2 after treating NB WT mice with CSF1 antibody or control IgG at NB. The cell amount of pitMØ, Ly6C^Low^MO and Ly6C^Hi^MO cells in the WT mouse pituitary gland at P2 after CSF1 antibody or control IgG treatment at NB. The quantitative data are shown as mean ± SEM. Each dot represents data from mice pooled together NB (n=9-10 samples/pool). Data are from 5 individual experiments. (F) Experimental outline for studying pituitary macrophages and monocytes at P2 after treating NB WT mice with CDF1R antibody or control IgG at NB. The cell amount of pitMØ, Ly6C^Low^MO and Ly6C^Hi^MO cells in the WT mouse pituitary gland at P2 after CSF1R antibody or control IgG treatment at NB. The quantitative data are shown as mean ± SEM. Each dot represents data from mice pooled together NB (n=7-11 samples/pool). Data are from 5 individual experiments.

Following the scRNA-seq data, we then divided the pitMØ into the pBAM-MAC population, identified by the expression of CD206, and into the ml-MAC subpopulation based on the expression of P2RY12 (Figure S2A). The similarity in the rest of the macrophage clusters in transcriptomics (Core-MACs, MHC II^Pos^-MACs, Oxt-MACs, and Inf-MACs) prevented us from distinguishing them individually by flow cytometry. However, at the transcriptomics level, these cells were divided by MHC II expression; thus, we collectively referred to those populations as MHC II^Pos^-MAC or MHC II^Neg^-MAC populations (Figure S2A). At E17.5, the CD206-positive pBAM-MACs formed around 35 % of pitMØ in the developing pituitary gland. Onward, the pBAM-MAC macrophage population diminished after 1 week of age (Figure 2B), as observed in scRNA-seq data (Figures 1D and S1E). Likewise, the P2RY12-positive ml-MACs were already present in the E17.5 pituitary gland (Figure 2B). Contradictory to the pBAMs, the ml-MAC population remained detectable over time (Figure 2B). During the early development of the pituitary gland (E17.5 and NB), only MHC II^Neg^-MACs were found. However, the expression of MHC II increased significantly in the postnatal pituitary gland (Figure 2B). Interestingly, both the disappearance of pBAMs and the appearance of MHC II^Pos^-MACs were evident during the same period from 1-to 2 weeks. This simultaneous shift suggests a potential alteration in macrophage functionality, particularly in their antigen-presenting role to CD4^Pos^ T cells within the pituitary gland (Figure S1A-S1C). In immunostaining of macrophages with F4/80, we observed that pitMØ were scattered throughout the pituitary gland stroma, both in anterior and posterior lobes, but, as shown previously ^33,43^, were not frequent in the intermediate lobe of the pituitary gland (Figures 2C, S2D and S2E). At early time points (E17.5, NB, and 1 week), many macrophages exhibited a more rounded phenotype, whereas, in the pituitaries of 2-to 8-week-old mice, macrophages presented the typical morphology of mature phagocytes with multiple dendrites. F4/80 positive cells were localized both connected to vessels and within the pituitary gland stroma (Figure 2D).

The colony-stimulating factor 1 receptor (CSF1R, also known as M-CSFR or CD115) is essential for macrophage development, differentiation, and survival ^44,45^. The receptor has two ligands, colony-stimulating factor 1, CSF1, and interleukin (IL)-34 ^46^. To assess if CSF1 or IL-34 is required to maintain the pitMØ populations, we first used IL-34-deficient reporter mice ^47^. We found that IL-34 was predominantly expressed in the anterior pituitary gland (Figure S2F). However, unlike the brain of IL-34 deficient mice displaying a decrease in macrophage numbers ^46,47^, the pitMØ were present at normal numbers in IL-34 deficient mice (Figures S2G), suggesting that IL-34 has no role in the development or maintenance of the pitMØ. We next gave a single injection of function-blocking CSF1 antibody (clone 5A1), preventing the binding to CSF1R, or a neutralizing antibody CSF1R (clone AFS98) for NB mice (Figures 2E and 2F). Both antibodies were sufficient to cause a significant reduction of pitMØ 48h after treatment (Figures 2E and F). Notably, depletion was evident equally in different macrophage subpopulations (Figure S2H). Both monocyte subsets, Ly6^Hi^MO and Ly6C^Low^MO, were unaffected by CSF1 and CSF1R antibody injections (Figure 2E and 2F). Hence, the result confirmed that CSF1R signaling *via* CSF1 is critical for pitMØ homeostasis.

### The pituitary gland macrophages are located in unique spatial niches and have close interaction with hormone-secreting cells

We next examined whether different pitMØ subpopulations have a specific microanatomical niche in the pituitary gland. In the developing pituitary gland, CD206^Pos^ pBAM-MACs were predominantly located in the anterior pituitary gland (Figures 3A and S3A). The pBAM-MACs also showed LYVE-1 positivity (Figure S3B). The pBAM-MAC cells were closely mingling with blood vessels (Figure 3A, S3A, and S3B), in line with to previous studies displaying CD206 and LYVE-1 as markers of vascular association ^13,38^. Similarly to flow cytometry results, only a few CD206-expressing pBAM-MACs were found in the pituitary gland of the older animals (5-week-old). In contrast to pituitaries of E17.5 and NB animals, the few remaining pBAM-MACs in 5-week-old mice were mainly located in the pituitary gland stalk area, connecting the pituitary gland to the hypothalamus, where the superior hypophyseal artery and portal veins are located (Figure 3B). MHC II^Pos^-MACs were only detectable in the pituitary gland of 2-week-old and older mice (Figures 2B and 3C). MHC II^Pos^-MACs were situated mainly in the anterior pituitary gland, with no detection in the posterior lobe. However, we found a small number of MHC II positive cells that resemble the characteristics of macrophages on the boundary of the intermediate lobe (Figure 3C). Notably, the ml-MACs, positive for P2RY12, were solely localized to the stroma of the posterior lobe in all timepoints studied (Figure 3B and 3C). Interestingly, ml-MACs showed highly branched morphology similar to brain microglia ^48,49^ and were not connected to blood vessels (Figures 3C and S3C). We also observed a more roundish shape, with only F4/80 positive macrophages, which were mainly detected close to blood vessels (Figure S3C). We validated our observation by flow cytometry by separating the anterior and posterior lobes of the pituitary gland and were able to assign the P2RY12 positive ml-MACs only to the posterior lobe of the pituitary gland (Figure S3D).

**Figure 3.**
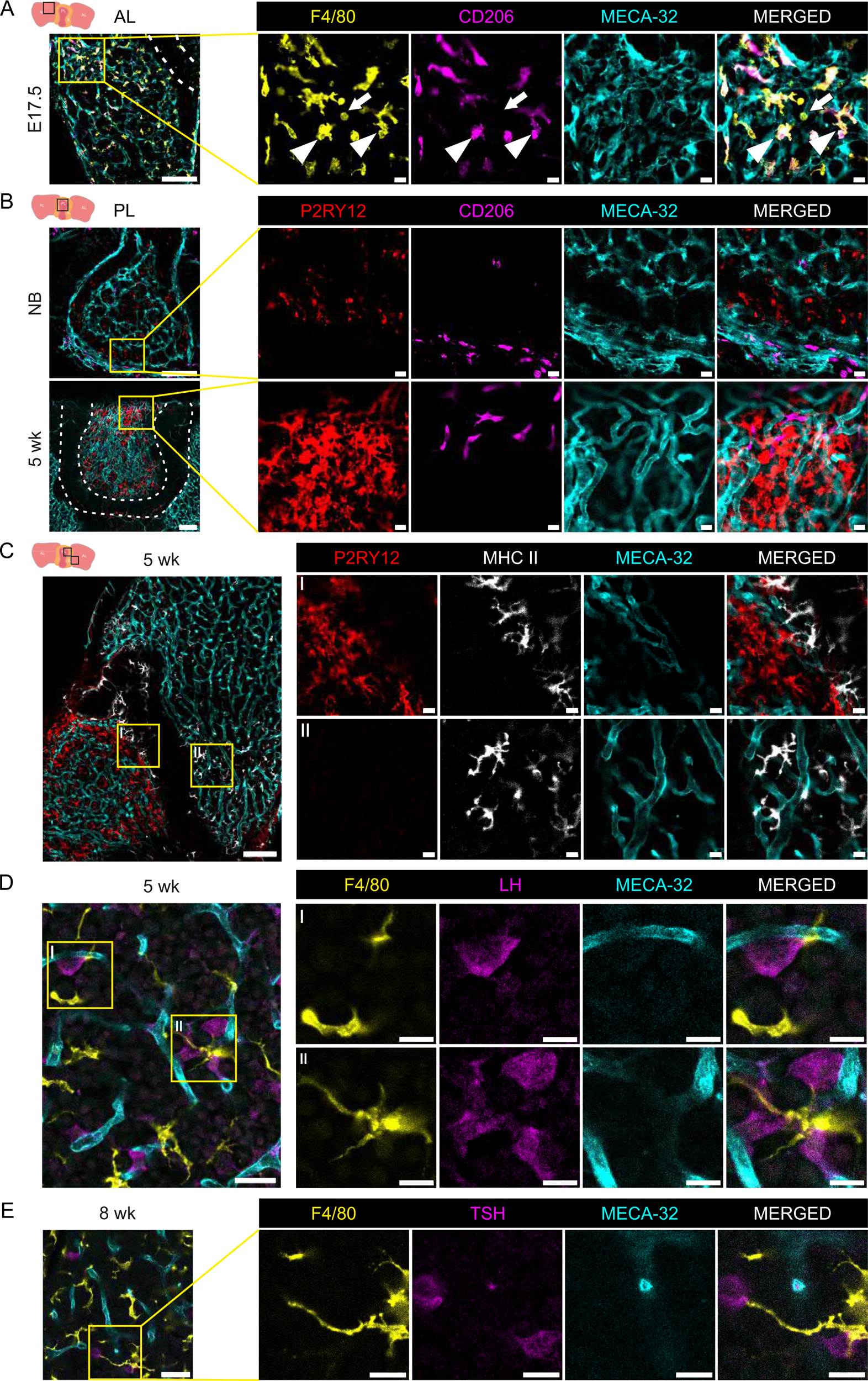
The pituitary gland macrophages are located in unique spatial niches and have close interaction with hormone-secreting cells. (A) Pituitary gland whole mount immunofluorescence with F4/80, CD206 and endothelial marker, MECA-32, from male WT pituitary glands at E17.5. Inserts are higher magnifications from the boxed anterior lobe area. Arrows indicate F4/80^Pos^CD206^Neg^ cells and arrowheads indicate F4/80^Pos^CD206^Pos^. For overview image, scale is 100 µm, and for inserts, 10 µm. Shown are representative images from 3 independent experiments. (B) Pituitary gland whole mount immunofluorescence with P2RY12, CD206 and endothelial marker, MECA-32, from male WT pituitary glands at NB and 5 weeks of age. Inserts are higher magnifications from the boxed posterior lobe area. For overview images, scales are 100 µm, and for inserts, 10 µm. Shown are representative images from ≤ 3 independent experiments. (C) Pituitary gland whole mount immunofluorescence with P2RY12, MHC II and endothelial marker, MECA-32, from male WT pituitary gland of 5 weeks of age. Inserts are higher magnifications from the boxed posterior lobe area (I) and the boxed anterior lobe area (II). For overview image, the scale is 100 µm, and for inserts, 10 µm. Shown are representative images from 3 independent experiments. (D) Pituitary gland whole mount immunofluorescence with F4/80, luteinizing hormone (LH) and endothelial marker, MECA-32, from male WT pituitary gland of 5 weeks of age. Inserts are higher magnifications from the boxed anterior areas. For overview image, the scale is 100 µm, and for inserts, 10 µm. Shown are representative images from 3 independent experiments. (E) Pituitary gland whole mount immunofluorescence with F4/80, thyroid-stimulating hormone (TSH) and endothelial marker, MECA-32, from WT pituitary gland of 8 weeks of age. Inserts are higher magnifications from the boxed anterior area. For overview image, the scale is 100 µm, and for inserts, 10 µm. Shown are representative images from 3 independent experiment.

We next investigated the spatial relationship between macrophages and endocrine cells in the anterior pituitary by conducting whole mount imaging analyses using markers for macrophages (F4/80) and luteinizing hormone (LH, associated with gonadotrophs) or thyroid-stimulating hormone (TSH, associated with thyrotrophs) (Figures 3D and E). As previously observed ^50,51^, gonadotrophs were closely associated with blood vessels. Intriguingly, macrophages were also observed in close proximity to gonadotrophs, primarily making contact with them through their protrusions that surrounded the gonadotrophs (Figure 3D). Macrophages were also observed to be closely associated with thyrotrophs. However, unlike the interaction with gonadotrophs, macrophages appeared to exhibit protrusions that connected nearby thyrotrophs, indicating a distinct mode of interaction compared to their association with gonadotrophs (Figure 3E and S3E). Altogether, our findings unveil a particular spatial distribution of phenotypically diverse macrophages within the pituitary gland and suggest potential interactions and communication between macrophages and hormone-secreting cells in the microenvironment.

### Pituitary gland macrophages originate from early yolk sac progenitors

The ontogeny of pituitary macrophages (pitMØ) has not been studied. Therefore, we employed a combination of established macrophage reporter mice to unravel the origin and persistence of tissue-resident pitMØ. In the brain, microglia and BAMs derive from yolk sac-origin macrophage progenitors, and in homeostatic conditions, neither of the populations is replaced by monocytes from the bone marrow ^13,37,52^. Based on the observed similarities of pitMØ to the brain macrophages, we first generated reporter animals by crossing *Cx3cr1^CreERT^*^2^ mice ^53^ to *R26^EYFP^* or *R26^tdTomato^*mice ^54^ to tag the yolk sac macrophages and yolk sac erythro-myeloid precursor (EMP) origin macrophages. Pregnant dams were injected with tamoxifen (TAM) at E7.5, E8.5, or E9.5 to induce irreversible reporter expression in yolk sac primitive macrophages (TAM E7.5) or the EMPs and their progeny (TAM E8.5 or E9.5). As Cre-induced excision is not 100 % penetrant, we used the labeling of microglia as a control for the efficiency of pulse-labeling experiments. Consistent with a previous report ^52^, analyses of the brains of E17.5 embryos pulse labeled at E7.5, E8.5, or E9.5 showed high labeling of microglia in the brain (Figure S4A, S4B, and S4C).

In the pituitary gland, whole mount analysis at E17.5, pulse labeling already at E7.5 revealed a distinctive presence of tdTomato-positive macrophages, signifying the contribution of primitive YS macrophages to the pitMØ (Figure 4A). Subsequently, the administration of TAM at E8.5 resulted in labeling tdTomato-positive macrophages in the pituitary gland at E17.5 (Figure 4B). Notably, both pulse labeling time-points led to the distribution of tdTomato-positive macrophages throughout the anterior and posterior regions of the pituitary gland (Figures 4A and 4B). Furthermore, initiation of labeling at E9.5 demonstrated EYFP-positive cells exclusively within the pitMØ population, with both Ly6C^Low^MO and Ly6C^Hi^MO evading the labeling (Figure 4C and S4D). Notably, EYFP labeling was observed both in CD206^Pos^ (pBAM-MACs) and CD206^Neg^ (including ml-MACs and MHC II^Neg^-MAC) cells (Figure S4E).

**Figure 4.**
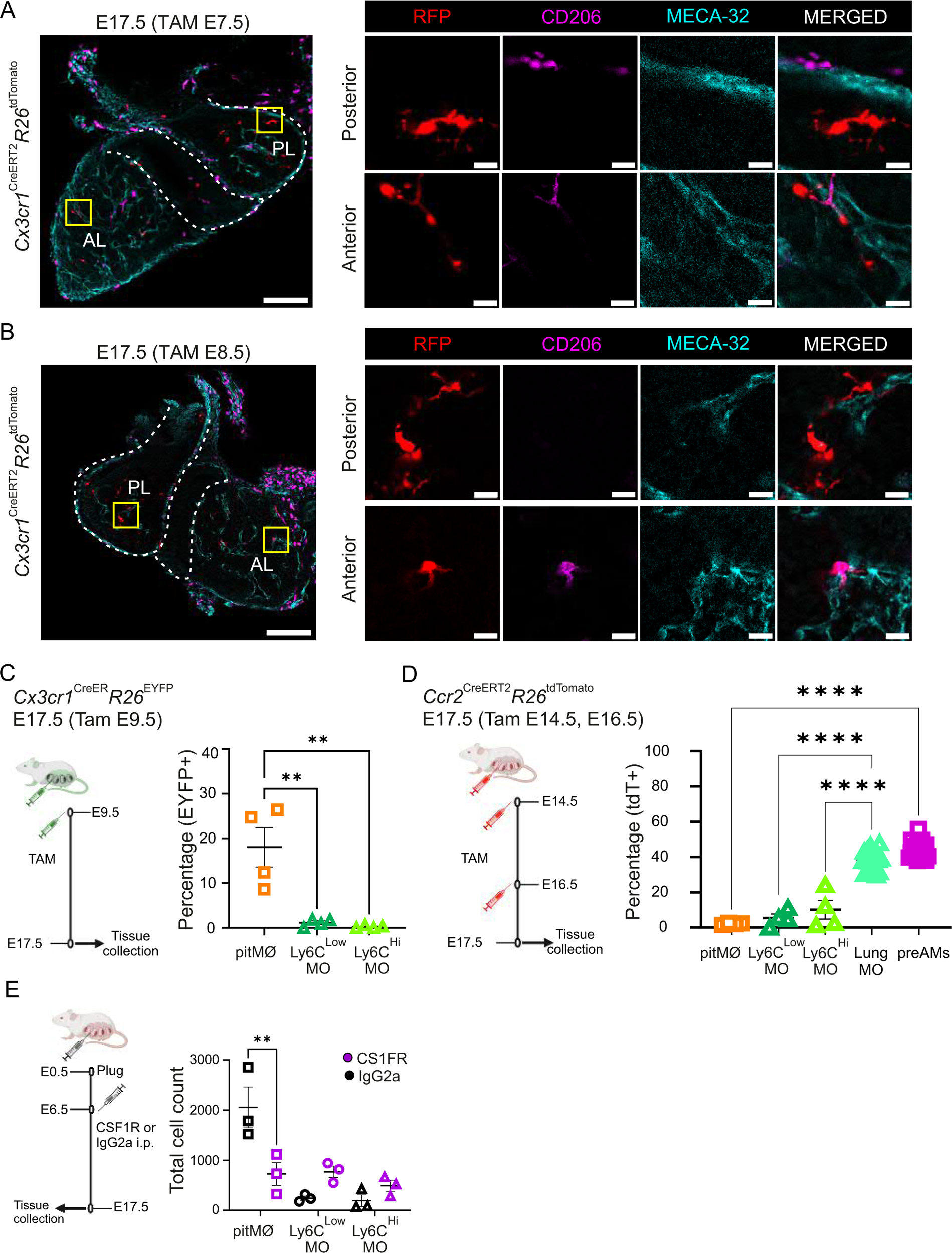
Pituitary gland macrophages originate from early yolk sac progenitors. (A, B) Pituitary gland whole mount immunofluorescence with RFP, CD206 and endothelial marker, MECA-32, from E7.5 (A) or E8.5 (B) tamoxifen labelled Cx3cr1^CreERT2^R26^tdTomato^ pituitary glands at E17.5. Inserts are higher magnifications from the boxed posterior lobe (PL) and anterior lobe (AL) area. For overview images, scales are 100 µm, and for inserts, 10 µm. Shown are representative images from 3 independent experiments. (C) Experimental outline and frequencies of EYFP expression in pitMØ, Ly6C^Low^MO and Ly6C^Hi^MO populations of E9.5 tamoxifen labelled Cx3cr1^CreERT2^R26^EYFP^ pituitary glands at E17.5. The quantitative data are shown as mean ± SEM. Each dot represents data from mice pooled together (n=9-11 samples/pool). Data are from 4 individual experiments. (D) Experimental outline and frequencies of tdTomato expression in pitMØ, Ly6C^Low^MO, Ly6C^Hi^MO, lung monocytes (Lung MO) and pre-alveolar macrophages (preAMs) of E14.5 and E16.5 tamoxifen labelled Ccr2^CreERT2^R26^tdTomato^ mice at E17.5. The quantitative data are shown as mean ± SEM. Each dot represents data from mice pooled together (pitMØ, Ly6C^Low^MO, Ly6C^Hi^MO, n=8-10 samples/pool) or one animal (Lung MO, preAMs, n=19). Data are from 4 individual experiments. (E) Experimental outline for studying pitMØ and monocytes at E17.5 after treating WT embryos with blocking CSF1R antibody or control IgG at E6.5. The cell amount of E17.5 pitMØ, Ly6C^Low^MO and Ly6C^Hi^MO cells in the WT mouse pituitary gland after CSF1R antibody or control IgG treatment. The quantitative data are shown as mean ± SEM. Each dot represents data from mice pooled together (n=8-11 samples/pool). Data are from 3 individual experiments.

The yolk sac-origin late EMPs not only give rise to non-monocytic yolk sac macrophages ^55^ but also migrate to the fetal liver to generate the *Cx3cr1*-expressing fetal liver monocytes ^56^. To evaluate embryonic liver monocyte involvement in the pitMØ, we used *Ccr2^CreERT^*^2^*; R26^tdTomato^* reporter mice, which allows the tracking of the Ly6C^Hi^ CCR2^+^ fetal liver-derived monocytes ^13,57^. Pregnant *Ccr2^CreERT^*^2^*; R26^tdTomato^* mice were treated with tamoxifen at E14.5 and E16.5. As expected, fetal monocytes in blood, liver, and lung, and monocyte-origin premature alveolar macrophages (preAMs) were tdTomato positive (Figures 4D, S4F-J). A low frequency of Ly6C^Hi^MO cells was labeled in the pituitary gland (Figure 4D). In contrast, pitMØ did not show any labeling, confirming that circulating CCR2-positive fetal liver monocytes do not engraft in the pituitary gland (Figure 4D). To further selectively cross-examine the putative yolk sac-origin of the pitMØ, we injected a single dose of function-blocking anti-CSF1R (AFS98) antibody into WT pregnant dams at E6.5 to selectively ablate yolk sac-origin macrophages from the developing embryos. The flow cytometric analyses of E17.5 pituitary glands verified the significant depletion of pitMØ while monocytes remained at the usual level (Figure 4E). Altogether, data suggest that pitMØ originate from yolk sac macrophages and yolk sac EMP descendants, similar to brain microglia and BAMs.

### The pituitary gland macrophages are maintained with self-renewing without an input of bone marrow-derived monocytes

Tissue-resident macrophages are maintained in tissues by local self-proliferation or monocyte recruitment from blood circulation ^15,58,59^. We performed whole mount immunostaining with anti-Ki67 and anti-F4/80 antibodies to characterize the macrophage proliferation in the developing pituitary gland. During postnatal development, the pituitary gland undergoes significant tissue expansion driven by the proliferation of differentiated endocrine cells and stem/progenitor cells in response to proliferative signals. Accordingly, our whole mount Ki67 staining revealed proliferating cells scattered throughout the pituitary gland of 1-week-old mice (Figure 5A). Upon closer examination, we observed proliferating macrophages in both the anterior and posterior areas of the pituitary gland, suggesting that pitMØ underwent *in-situ* proliferation (Figure 5A and S5A). This finding indicated that self-renewal through proliferation plays a role in maintaining the population of pituitary macrophages under steady-state conditions. However, in many postnatal tissues, embryonic-derived tissue-resident macrophages are entirely or partially replenished by bone marrow monocytes recruited from the blood circulation ^59–61^. Therefore, we next sought to determine the replenishment of embryonic origin cells and the contribution of adult bone marrow-derived monocytes in the pituitary gland by using *Ms4a3^creERT^*^2^*; R26^tdTomato^* monocyte fate-mapping reporter mice ^62^, which allows distinguishing between embryonically seeded (tdTomato^Neg^) and adult monocyte-derived (tdTomato^Pos^) macrophages. Notably, pitMØ do not intrinsically express *Ms4a3* and could attribute the tdTomato labeling only after tamoxifen-induction through monocyte recruitment. At 4 weeks of age, *Ms4a3^creERT^*^2^*; R26^tdTomato^* reporter mice were given intraperitoneal (i.p.) tamoxifen injection for 5 consecutive days, and we first analyzed the pituitary gland and blood after 4 days from the last injection (Figure 5B). As anticipated, blood monocytes were efficiently tagged (Figures S5B and S5C). However, within the pituitary gland, tdTomato positivity was almost exclusively observed in the monocyte populations, while the pitMØ exhibited no tdTomato positivity, indicating that their development proceeded autonomously, unaffected by the influence of bone marrow monocytes (Figure 5C).

**Figure 5.**
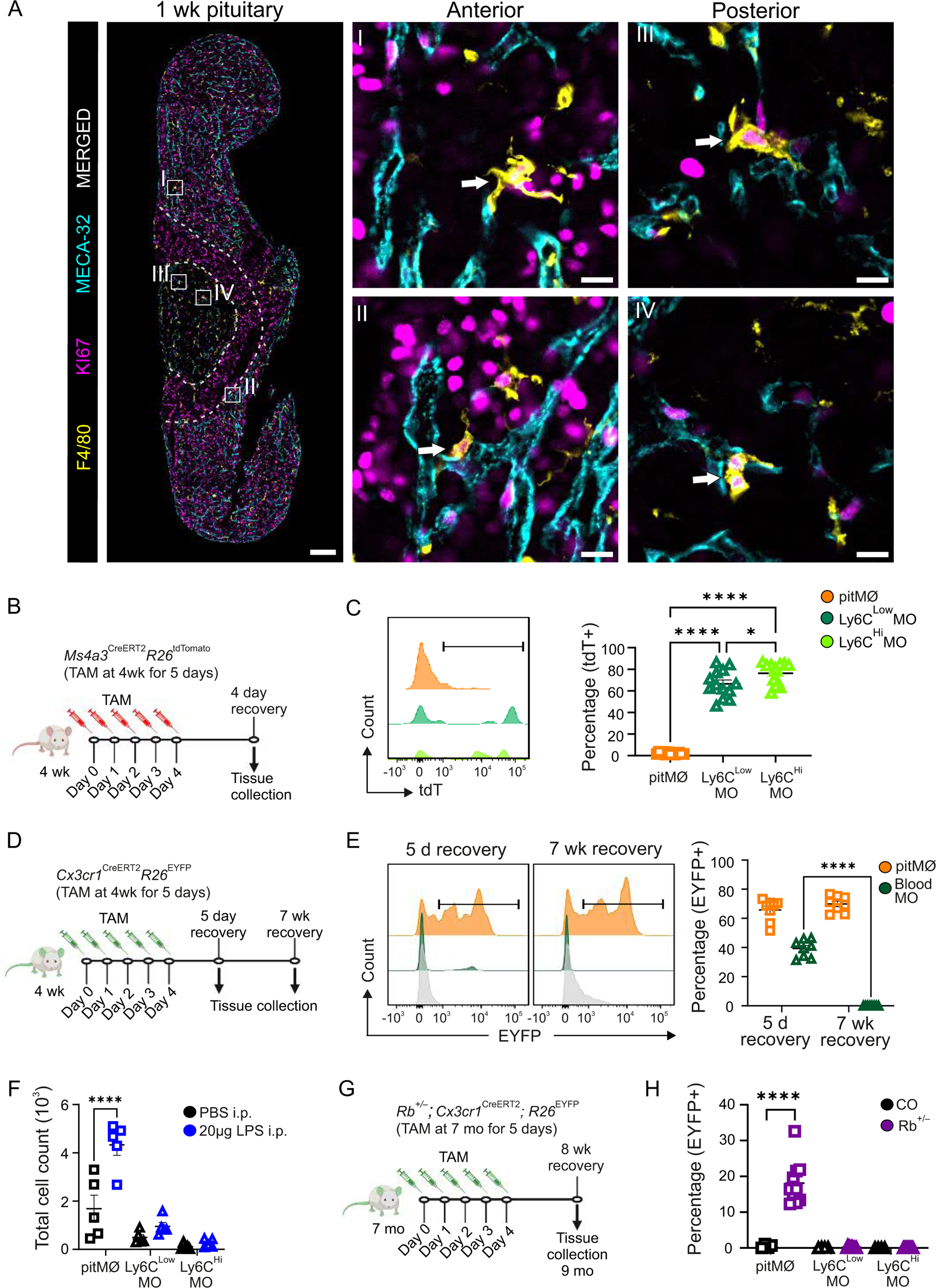
Early yolk sac-derived pituitary gland macrophages self-maintain until late adulthood by proliferation. (A) Pituitary gland whole mount immunofluorescence with F4/80, proliferation marker KI67, and endothelial marker, MECA-32, from WT pituitary glands of 1-week-old mice. Inserts are higher magnifications from the boxed anterior lobe area (I-II) and the boxed posterior lobe area (III-IV). Arrows indicate F4/80^Pos^KI67^Pos^ cells. For overview images, scales are 100 µm, and for inserts, 10 µm. Shown are representative images from 3 independent experiments. (B, C) Experimental outline, representative histograms and frequency of tdTomato^Pos^ cells in pitMØ, Ly6C^Low^MO and Ly6C^Hi^MO cells in the Ms4a3^CreERT2^R26^tdTomato^ mouse pituitary gland at 5 weeks of age (4 day recovery). Tamoxifen induction for five consecutive days. The quantitative data are shown as mean ± SEM. Each dot represents data from one mouse (n=13). Data are from 4 individual experiments. (D, E) Experimental outline, representative histograms and frequency of EYFP^Pos^ cells in pitMØ and blood monocytes in the Cx3cr1^CreERT2^R26E^YFP^ mouse pituitary gland at 5 weeks of age (5 day recovery) and 12 weeks of age (7 wk recovery). Tamoxifen induction for five consecutive days. The quantitative data are shown as mean ± SEM. Each dot represents data from one mouse (n=8). Data are from 3 independent experiments. (F) Cell amount of pitMØ, Ly6C^Low^MO and Ly6C^Hi^MO cells in the WT mouse pituitary gland at 10 weeks of age after LPS injection (i.p.) and 24 hour recovery. The quantitative data are shown as mean ± SEM. Each dot represents data from one mouse (n=5). Data are from 3 independent experiments. (G, H) Experimental outline, cell amount and frequency of EYFP^Pos^ cells in pitMØ, Ly6C^Low^MO and Ly6C^Hi^MO in the Cx3cr1^CreERT2^R26E^YFP^;Rb^-/+^ mouse pituitary gland compared with littermate controls at 9 months of age. Tamoxifen induction for five consecutive days at 7 months of age. The quantitative data are shown as mean ± SEM. Each dot represents data from one mouse (n=4-10). Data are from 3 independent experiments.

As an alternative way to explore the replenishment of the pitMØ, we used the *Cx3cr1^CreERT^*^2^*; R26^EYFP^* model. At the age of 4 weeks, *Cx3cr1^CreERT^*^2^*; R26^EYFP^* animals underwent a regimen of tamoxifen administration for five consecutive days. This approach was employed to ensure a robust and irreversible induction of EYFP in all CX3CR1-positive cells, including blood monocytes (Figure 5D). Over time, cells developing from blood monocytes after tamoxifen withdrawal are no longer labeled, while long-lived or self-renewing cells will remain EYFP-positive. We collected tissues 5 days or 7 weeks after the final tamoxifen dose to assess pitMØ replenishment by blood monocytes. As shown previously, blood monocytes were EYFP^+^ 5 days after the last tamoxifen dose before being replaced by EYFP^−^ monocytes during the 7 weeks of recovery time (Figure 5E) ^53,63^. CX3CR1^+^ microglia, which do not undergo replenishment by blood monocytes, consistently retained a high level of EYFP expression even after a 7-week recovery period (Figure S5D and S5E). In the pituitary gland, 5 days post-TAM injection, 65.8 % ±3 of pitMØ were EYFP positive (Figure 5E). Remarkably, even after 7 weeks of recovery, the labeling of pituitary gland pitMØ consistently maintained EYFP positivity, indicating that these macrophages were locally maintained through proliferation (Figure 5E). Further supporting this notion, a single TAM pulse administered at E16.5 resulted in the persistence of EYFP-tagged macrophages at 7 and 14 weeks of age (Figure S5F). Furthermore, we quantified macrophages in CCR2-deficient mice, which have severely reduced blood monocyte counts^64^. Despite the reduced monocyte number in blood, the macrophage counts in the pituitary gland were not decreased in CCR2-deficient mice, supporting the role of local proliferation in maintaining the pitMØ pool (Figure S5G). To study whether bone marrow-derived monocytes can infiltrate the pituitary gland in pathophysiological conditions, we stimulated acute inflammation by i.p. lipopolysaccharide (LPS) injection to adult WT mice. Flow cytometry analysis of the pituitary glands and serum IL-6 measurement were performed 24 hours after LPS injection. Interestingly, the infection caused an accumulation in pitMØ but did not increase monocyte numbers within the pituitary gland (Figure 5F), despite observing a significant increment in serum IL-6 levels after 24 hours (Figure S5H). These results indicate that, even during systemic inflammation, bone marrow-derived monocytes are not recruited into the pituitary gland.

Pituitary gland adenomas present the most common disease affecting the pituitary gland ^65^. We utilized heterozygous tumor suppressor Retinoblastoma (Rb^+/-^) deficient mice to investigate how pituitary gland adenomas influence the pitMØ. These mice are shown to develop pituitary gland adenomas with 100 % penetrance in old mice ^66,67^. To specifically examine the impact on pitMØ, the *Rb^+/-^* mice were crossed with *Cx3cr1^CreERT^*^2^*; R26^EYFP^* mice. Subsequently, 7-month-old *Rb^+/-^*; *Cx3cr1^CreERT^*^2^*; R26^EYFP^*mice were treated with tamoxifen for 5 consecutive days and analyzed after 8-week recovery time (at the age of 9 months; Figure 5G). Interestingly, the presence of pituitary gland adenomas did not increase in the total numbers of pitMØ or monocytes in *Rb^+/-^*; *Cx3cr1^CreERT^*^2^*; R26^EYFP^*mice (Figures S5I and S5J). Notably, even after a substantial time after the last tamoxifen injection, the pitMØ in *Rb^+/-^*; *Cx3cr1^CreERT^*^2^*; R26^EYFP^* mice remained EYFP positive (Figure 5H). In summary, these findings indicate that pitMØ are maintained without input from bone marrow-derived circulating monocytes in adult animals, even in the presence of pituitary gland disorders such as adenomas.

### The presence of early macrophages plays a crucial role in maintaining hormonal equilibrium within the pituitary gland

We next addressed the functional importance of macrophages for normal pituitary gland development and function. We ablated the pre-existing tissue-resident macrophages with function-blocking anti-CSF1R antibody. We gave the first injection during embryo development at E6.5 to block the early seeding of yolk sac macrophages in the pituitary gland. After birth, an anti-CSF1R antibody was given every second week (4 times in total), and tissues were collected one week after the last injection (at the age of 5 weeks; Figure 6A). The depletion of macrophages did not yield a noteworthy difference in weight gain between the control and anti-CSF1R-treated mice (Control 17.81 ± 0.45 g versus 16.47 ± 0.77 g; n = 11-16). However, the treatment efficiently and persistently depleted macrophages in the pituitary gland (Figure S6A). At the same time, the numbers of brain microglia in anti-CSF1R-treated mice were comparable to those in the control group (Figure S6B).

**Figure 6.**
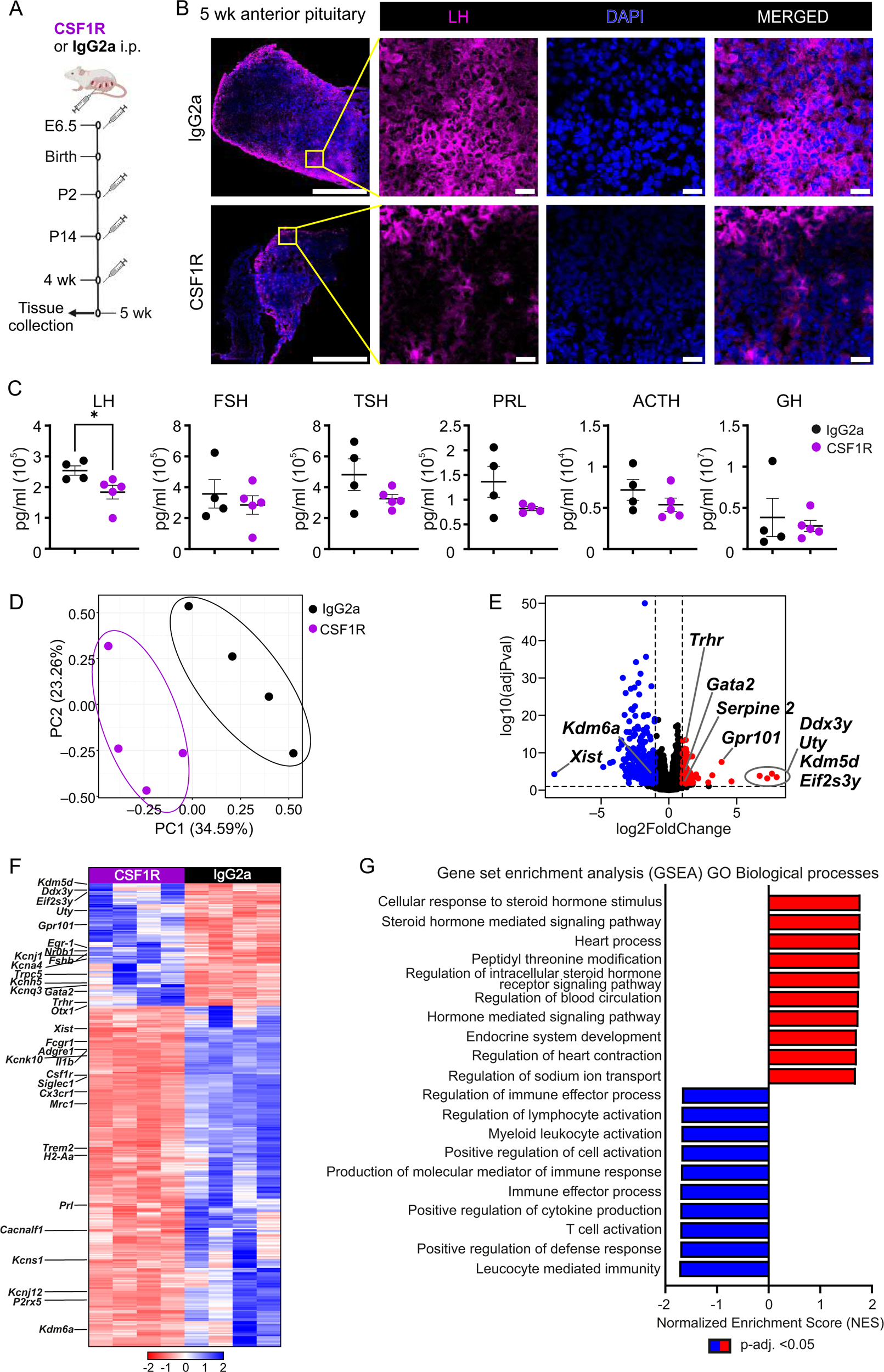
The presence of early macrophages plays a crucial role in maintaining hormonal equilibrium within the postnatal pituitary gland. (A) Experimental outline for studying pitMØ and monocytes of 5-week-old WT mice after treated with blocking CSF1R antibody or control IgG at E6.5, postnatal day (P) 2, P14 and 4 weeks of age. (B) Pituitary gland frozen section immunofluorescence staining with anti-LHβ and nuclear marker DAPI from pituitary gland of 5-week-old WT mice after treated with blocking CSF1R antibody or control IgG at E6.5, P2, P14 and 4 weeks of age. Inserts are higher magnifications from the boxed anterior lobe areas. For overview image, scale is 500 µm, and for inserts, 20 µm. Shown are representative images from 3 independent experiments. (C) Intra-pituitary hormone measurements for LH, FSH, TSH, PRL, ACTH and GH from pituitary gland of 5-week-old WT mice after treated with blocking CSF1R antibody or control IgG at E6.5, P2, P 14 and 4 weeks of age. The quantitative data are shown as mean ± SEM. Each dot represents data from individual mice (n=4-5). (D) Principal Component Analysis (PCA) of the bulk RNA-seq data from pituitary gland of 5-week-old WT mice after treated with blocking CSF1R antibody or control IgG at E6.5, P2, P14 and 4 weeks of age. Data from individual mice (n=4). (E-F) Volcano plot (E) and heatmap (F) of the Differentially Expressed Genes (DEGs) between the pituitary glands of 5-week-old WT mice after treated with blocking CSF1R antibody or control IgG at E6.5, P2, P 14 and 4 weeks of age. Data from individual mice (n=4). (G) Gene set enrichment analysis (GSEA; GO Biological processes) of the bulk RNA-seq data. Gene sets are ranked according to their normalized enrichment scores (NES).

The primary function of the pituitary gland is hormone production and secretion. To investigate the impact of macrophage depletion on this crucial role, we conducted an immunohistochemical analysis focusing on cells responsible for producing LH, prolactin (PRL), and TSH associated with gonadotrophs, lactotrophs, and thyrotrophs, respectively (Figures 6B and S6C-E). Examination of the pituitary glands did not reveal apparent disparities in the proportions of hormone-secreting cell populations in the macrophage-depleted pituitaries (Figures 6B, S6D, and S6E). However, we noted a reduced intensity of LH and TSH staining in macrophage-depleted pituitaries compared to controls (Figures 6B and S6D), while the intensity of PRL staining remained unchanged (Figure S6E). Additionally, the spatial organization of LH-producing gonadotrope cells appeared disorganized in macrophage-depleted pituitaries (Figure 6B).

We next investigated whether the identified alterations exert a functional impact on the hormonal milieu of the pituitary gland in macrophage-depleted mice. To assess this, we measured intrapituitary hormonal levels. Intriguingly, among all six hormones produced by the anterior pituitary, we observed a significant reduction in intrapituitary luteinizing hormone (LH) concentration (Figure 6C). Conversely, no significant decrease was measured in pituitary follicle-stimulating hormone (FSH) despite both hormones being produced by gonadotrope cells (Figure 6C). Furthermore, reductions in TSH and PRL hormone levels were observed, although these changes did not reach statistical significance. In contrast, levels of adrenocorticotropic (ACTH) and growth (GH) hormones were similar between treated and control mice (Figure 6E and 6C), suggesting a hormone-secreting cell-specific role of macrophages in the pituitary gland. We then limited the depletion of macrophages exclusively to the embryonic developmental stage. Remarkably, administering the anti-CSF1R antibody only once at E6.5 also led to a compelling trend towards reduced intrapituitary hormonal levels at 4 weeks of age (Figure S7A). A pivotal regulator of LH and FSH synthesis and secretion is the hypothalamus-released gonadotropin-releasing hormone (GnRH). Gonadotrope cells respond to GnRH through the specific expression of the GnRH receptor (*Gnrhr*). To eliminate the possibility of extended macrophage depletion leading to a reduction in hormone levels as a secondary consequence of its impact on hypothalamus GnRH release, we analyzed the *Gnrhr* expression in macrophage-depleted pituitaries. Importantly, we noticed that *Gnrhr* expression was not compromised in macrophage-depleted pituitaries (Figure S7B), indicating that the observed reduction in LH levels was not attributed due to the deficiency in GnRH signaling. Collectively, these findings point towards an impaired hormonal milieu within the anterior pituitary gland of macrophage-deficient mice and suggest that tissue-resident macrophages play a role in the intricate regulation of hormonal balance within the pituitary gland.

To further gain insights into the molecular signature of macrophage-depleted pituitary glands, we performed bulk RNA sequencing (bulk RNA-seq) of whole pituitary glands of the anti-CSF1R and control-treated WT mice. Principal component analysis revealed a clear difference between anti-CSF1R and control-treated pituitary glands (Figure 6D), having 1438 differentially expressed genes (DEGs) altogether (Table S2). Most of the DEGs (74 %; Fold change >1,6 adjusted p-value <0.1) were downregulated in macrophage-depleted pituitary gland compared with the control. Among the downregulated genes, a substantial number were core macrophage signature genes, including *Fcgr1*, *Adgre1*, *Mrc1*, *H2-Aa*, *Siglec1*, *Il1b*, *Cx3cr1*, *Trem2*, and *Csf1r* (Table S2). This robust suppression validates the effective depletion of macrophages. Interestingly, despite exclusively employing male mice in our experiment, male-biased Y chromosome-linked genes such as *Ddx3y*, *Uty*, *Kdm5d*, and *Eif2s3y* and several autosomal genes (e.g., *Drd4*) genes, also known to have a sex-biased expression ^68,69^ were upregulated following macrophage depletion. Simultaneously, female-biased genes like *Xist* and *Kdm6a*, associated with the X chromosome, exhibited downregulation (Figure 6E and Table S2).

Within the top DEGs, we identified several genes associated with hormone production in the anterior pituitary that exhibited upregulation in macrophage-depleted pituitary glands (Figures 6E and 6F and Table S2). These include the genes encoding pituitary hormone prolactin (*Prl*), thyrotropin-releasing hormone receptor (*Trhr*), follicle-stimulating hormone beta subunit (*Fshb*), and an orphan G-protein coupled receptor (*Gpr101*), which is involved in the growth of the pituitary gland and stimulus of GH ^70^. Moreover, expression of several transcription factors, including GATA-binding factor 2 (*Gata2*), crucial for gonadotrope and thyrotrope differentiation ^71^, nuclear receptor subfamily 0 group B member 1 (*Nr0b1*) involved in pituitary gonadotropins production ^72^, early growth response 1 (*Egr-1*) acting as a transcriptional mediator for GnRH-induced signals activating the LHβ gene ^73^, and Homeobox Protein (*Oxt1*) implicated in the juvenile stage control of GH, FSH, and LH hormone levels ^74^, was found to be enhanced in animals with depleted macrophages (Figure 6F and TableS2).

Endocrine cells display large-scale changes in coordinated calcium-spiking activity in response to various hypothalamic and peripheral stimuli. Elevation of cytosolic calcium (Ca^2+^) concentration is an essential trigger for hormone secretion and release in pituitary endocrine cells ^75^. These cells regulate Ca^2+^ levels through extracellular Ca^2+^ influx, accomplished with the mobilization of intracellular Ca^2+^ stores. Pituitary gland hormone-secreting cells express various voltage-gated sodium, calcium, potassium, and chloride channels and possess gap junction channels and extracellular ligand-gated ion channels, enabling extracellular Ca^2+^ influx ^76^. Interestingly, many genes related to Ca^2+^ influx were altered in macrophage-depleted pituitary (Figure 6F and Table S2). Expression of both short transient receptor potential channel 5 (*Trpc5*) and potassium voltage-gated channel subfamily Q member 3 (*Kcnq3*), pivotal in gonadotrope cells for inducing plasma membrane depolarization and enhancing intracellular Ca^2+^ levels upon GnRH stimulation ^77,78^, were amplified. Moreover, in macrophage-depleted animals, several other members of the voltage-gated channels exhibited upregulation, including *Kcnj1* (potassium inwardly rectifying channel subfamily J member 1), *Kcna4* (potassium voltage-gated channel subfamily A member 4), *Kcns1* (potassium voltage-gated channel modifier subfamily S member 1), *Kcnj12* (potassium inwardly rectifying channel subfamily J member) and *Cacng3* (calcium voltage-gated channel auxiliary subunit gamma 3) (Figure 6F and Table S2). Interestingly, a small G-protein Gem, an inhibitor of voltage-dependent calcium channels, also showed upregulation. On the other hand, conflicting results were also observed as several genes related to calcium signaling were downregulated, such as *Kcnk10* (potassium two-pore domain channel subfamily K member 10), *Kcnh5* (potassium voltage-gated channel subfamily H member 5), *Cacna1f* (calcium voltage-gated channel subunit alpha1 F) (Figure 6F and Table S2). Likewise, *P2rx5* (purinergic receptor P2X, ligand-gated ion channel, 5), which is primarily activated by extracellular adenosine triphosphate (ATP) and mainly found in LH and TSH cells in the anterior pituitary ^79^, was also down-regulated (Figure 6F and Table S2).

We next conducted gene set enrichment analysis analyses to understand how macrophage depletion affects molecular signaling and biological pathways in the pituitary gland. In our Gene Ontology (GO) analyses, we observed in anti-CSF1R-treated pituitary glands significant downregulation of immunological functions. These processes included T cell activation, response to bacteria, positive regulation of cytokine production, and regulation of innate immune response (Figure 6G). Conversely, the predominant up-regulated biological pathways were linked to hormonal regulation, encompassing cellular responses to steroids, steroid hormone-mediated signaling pathways, regulation of intracellular steroid hormone receptor signaling pathways, and endocrine system development (Figure 6G). Notably, among the top-regulated biological pathways were also many related to ion transport, such as the regulation of sodium ion transport and cardiac muscle contraction (Figure 6G). In addition, many of the top-up-regulated pathways identified using Reactome annotations were also related to ion transport (Figure S7C). To further explore the downstream functional consequences and to ensure the consistency of the bioinformatic analysis, we performed Ingenuity Pathways Analysis (IPA) on the differentially expressed genes. Furthermore, IPA pathway analyses (canonical pathways and biological functions) agreed with GSEA results and were linked to ion channel transport, potassium channels, and calcium signaling (Table S3). Together, the data indicates that macrophage depletion in pituitary glands profoundly impacts hormone production and suggests that macrophages have a role in the modulation of electrical activity and calcium influx in the pituitary gland.

## Discussion

Although macrophages have been reported in the pituitary gland ^32,33,43,80^, most of these studies have lacked sufficient resolution to profile the pituitary gland macrophage compartment comprehensively. In this study, we employed a combination of approaches to highlight the presence of transcriptionally and spatially variable resident macrophage subsets within the pituitary gland, showing dynamic phenotypic changes from neonatal to adulthood. Our results unveiled a distinct contrast in the transcriptional profiles of macrophage subpopulations between the anterior and posterior lobes and highlighted a notable divergence in their molecular characteristics. Fate-mapping analyses revealed that pitMØ originate solely from early yolk sac precursors and persists throughout the lifespan by local proliferation, even in pathophysiological conditions. Moreover, our findings demonstrated that depleting pitMØ has profound consequences on the hormonal balance of the pituitary gland.

The posterior pituitary lobe acts as a neuroendocrine interface connecting oxytocin and arginine-vasopressin axonal projections with the permeable capillary network of fenestrated endothelia. Our study unveiled a remarkable transcriptional and morphological similarity between the ml-MAC subset located in the posterior pituitary gland and microglia within the central nervous system (CNS) ^34–37^. Consequently, this observed likeness between ml-MACs and microglia can be attributed to the embryological and anatomical continuity between the posterior part of the pituitary gland and the hypothalamus. In contrast to neurovascular function in the posterior region, the anterior pituitary gland operates as a hormone-producing organ. Contrary to the homogeneity observed in macrophages within the posterior lobe, our results revealed that there is more transcriptional, kinetical, and spatial heterogeneity among the anterior lobe pitMØ than previously appreciated ^32,80,81^. During pituitary gland development, CD206^+^ vascular-associated pBAM-MAC was predominant in the anterior pituitary. However, as the pituitary gland matures, it constitutes only a minor fraction of the adult pitMØ compartment. Furthermore, in contrast to the ml-MACs found in the posterior pituitary, the majority of anterior pituitary macrophages exhibit a gradual increase in MHC II expression as animals mature, suggesting that macrophages are possibly preparing to acquire the ability to present antigens via MHC II molecules and produce cytokines, crucial for activating adaptive immune responses ^82^.

The initial wave of embryonic macrophages originates from early yolk sac myeloid precursors around E7.0. Following the establishment of the bloodstream around E9.5, primitive macrophages migrate into the developing tissues, and in the brain, they undergo differentiation to form microglia ^52,56^. In other tissues, early yolk sac macrophages are predominantly substituted in the later stages of development by monocyte-derived macrophages originating from the fetal liver ^83^. Post-birth, it is a common phenomenon that, over time, embryonic-derived macrophages are either entirely or partially displaced by CCR2-dependent Ly6C^Hi^ monocyte-derived macrophages ^59,84^. However, microglial cells stand out as distinctive exceptions among tissue-resident macrophages. In a state of homeostasis, these cells retain their unique yolk sac origin throughout adulthood ^52^. Utilizing advanced genetic lineage tracing, we show that, similarly to microglia ^13,38,52,85,86^, all the pitMØ, whether located in the posterior or anterior lobe, are seeded by the primitive yolk sac-derived macrophages and are not displaced by fetal liver monocytes-derived macrophages, although the pituitary gland is outside the blood-brain-barrier. Moreover, utilizing *Cx3cr1^CreERT^*^2^ ^53^ and *Ms4a3^CreERT^*^2^ ^62^ reporter mice, we found that renewal of the adult pituitary gland macrophage compartment during steady-state does not necessitate any contribution from bone marrow monocytes. In conclusion, our fate-mapping data suggest that, similar to microglia, the pituitary gland macrophages have a distinct origin exclusively from the yolk sac.

In a few tissues, such as the skin ^87^, lung alveolar space ^53,88^, and brain ^52^, only minor or no monocytes are observed under physiological conditions. This limited engraftment can be attributed to several factors, including the high self-renewal capacity of tissue-resident macrophages and restricted physical access due to barriers like the epithelium or the blood-brain barrier. However, in various disease states, including those affecting the brain, bone marrow-derived monocytes can infiltrate these tissues and differentiate into macrophages ^21^, and in the brain, BAMs have proven to be renewed through monocyte recruitment in adulthood and aging ^21,38^. Interestingly, despite the small and steady amount of monocytes in the pituitary and sustained macrophage depletion, monocyte-derived cells did not contribute to replenishing the empty macrophage niche in the pituitary gland. While microglial cells demonstrated re-population during the two-week recovery period following each antibody injection, pitMØ did not undergo replenishment within the same timeframe. It has been shown that the number of macrophages increases in the pituitary gland 6-24 h after LPS injection and returns to normal levels 48 h after ^89^. We observed the accumulation of pitMØ at 24 h when mice were subjected to acute LPS-induced inflammation, but even then, monocyte influx into the pituitary gland was not observed. The outcome was unexpected as it has been shown, employing a similar strategy, that systemic exposure to LPS induces the migration of inflammatory monocytes throughout various tissues, including the brain ^90^. Tumor-derived factors have been shown to attract circulating CCR2-expressing inflammatory monocytes into the tumor environment, where they differentiate into macrophages ^91,92^. Mice heterozygous for the retinoblastoma gene (*Rb*) develop ACTH-producing aggressive tumors arising from the intermediate lobe of the pituitary with high incidence from ages 2 to 11 months ^66,67^. Our studies reveal that tumor development in retinoblastoma-deficient mice did not increase the overall numbers of pitMØ or monocytes. Moreover, results from *Rb^+/-^*; *Cx3cr1^CreERT^*^2^; *R26^EYFP^* mice showed that, in agreement with the cell fate data, pitMØ populations showed no contribution from monocytes even with increasing age. Hence, our findings strongly suggest that pitMØ are maintained independently of bone marrow-derived monocytes, and even under pathophysiological conditions, increasing monocyte infiltration is not evident in the pituitary gland. However, it remains a possibility that prolonged LPS treatment or different types of pituitary tumors could result in increased monocyte migration within the pituitary gland, and future studies are essential to elucidate this aspect further.

The importance of the tissue microenvironment, inside which every cell interacts and is reciprocally influenced by its surroundings, has become clear in recent years. During pituitary development, secretory cells are not randomly spread throughout the pituitary gland but represent very organized three-dimensional network structures that play a functional role in integrating, amplifying, and transmitting signals ^6,93–95^. Our research suggests that prolonged depletion of macrophages disrupts the delicate networks of LH-producing gonadotrope cells and causes a decrease in intrapituitary LH levels. This finding is noteworthy because LH deficiency typically accompanies FSH deficiency due to their shared origin of secretion from gonadotroph cells ^3^. GnRH, produced in the hypothalamic neurosecretory cells and secreted in brief pulses from the hypothalamus, binds to GnRH receptors on the gonadotrope cells within the anterior pituitary and stimulates the synthesis and secretion of LH and FSH ^96^. Studies have revealed that the frequency of GnRH pulse plays a crucial role in determining the abundance of GNRHR on the cell surface ^97,98^. Moreover, it has been shown that null GNRHR mice exhibit low levels of FSH and LH and an incapacity to respond to exogenous GnRH ^99^. The systematic depletion of macrophages resulted in reduced levels of LH within the pituitary gland despite no apparent decrease in brain macrophage numbers, potentially leading to low GnRH levels. Importantly, there were no differences found in the expression of the *Gnrhr* between the macrophage-depleted or control pituitary glands, suggesting that the decrease in LH levels is not due to insufficient *GnRHR*-mediated signaling in pituitary gonadotropes.

Variations in gene expression between males and females are crucial in shaping sexual dimorphism and the physiological and behavioral differences between the sexes. This dimorphism is not only evident in external characteristics but also in the intricate molecular processes that regulate development and function. Research conducted in both rodents and humans has uncovered significant sex differences in the regulation of pituitary gland genes ^69,100^. Notably, these sex-specific distinctions in pituitary gene expression often manifest during and after puberty, a critical period characterized by increased hormonal synthesis and significant physiological changes ^2^. Surprisingly, our bulk RNA-seq, involving solely male mice, showed an increased expression of known male-biased genes such as *Ddx3y*, *Eif2s3y*, *Gpr101*, and *Drd4* ^69,101^. Conversely, genes associated with female traits, such as *Xist* and *Prl*, exhibited downregulation. These results suggest that male mice lacking macrophages have over-masculinized transcriptomes. This exciting discovery prompts questions about the adaptability of gene regulation in the absence of macrophages, emphasizing the need for additional research in future studies.

Regulating intracellular Ca^2+^ concentration is essential for hormone release. Pituitary gland hormone-secreting cells, like gonadotrophs, express a combination of numerous voltage-gated or dependent ion channels (e.g., calcium and potassium) to maintain calcium balance ^75,76,102^. Response to stimulus, such as a hypothalamic GnRH, leads to rapid fluctuations in intracellular Ca^2+^. The ion channels serve a dual purpose as they facilitate continuous calcium influx to sustain the plateau phase and, secondly, by replenishing the intracellular Ca^2+^ pools after hormone-induced calcium release. By coordinating these mechanisms, cells ensure appropriate responses to physiological stimuli while avoiding damaging consequences of Ca^2+^ dysregulation. After lifelong macrophage depletion, we observed multiple transcriptional alterations in genes associated with calcium homeostasis, such as *Cacng3* or *Trpc5*, which is predominantly expressed in juvenile mouse gonadotrophs and known to mediate calcium influx into cells upon activation by GnRH ^78^. Notably, we found that the expression of many genes related to depolarization and calcium influx was elevated, potentially affecting hormone synthesis and release.

ATP/ADP-dependent processes are shown to be crucial in balancing intracellular Ca^2+^ levels in various cells. This regulation is mediated through the activation of purinergic receptors, the G protein-coupled P2Y receptors (P2RYs), and ligand-gated P2X receptor-channels (P2XRs), by extracellular ATP or ADP binding. ATP/ADP provides a paracrine signaling mechanism for cross-talk between neighboring cells. Depletion of macrophages in the pituitary gland resulted in the downregulation of several genes coding purinergic receptors, like *P2ry6*, *P2rx7,* and *P2rx5*. In gonadotrophs, extracellular ATP amplifies GnRH-induced Ca^2+^ signaling and increases cytosolic Ca^2+^ by activating P2X2Rs. Reduction of extracellular Ca^2+^ has been shown to cause a prompt decline in LH release ^103–106^. The physiological sources of extracellular ATP responsible for activating purinergic receptors in the pituitary gland are not yet fully understood. However, it has been shown that macrophages can release ATP into the extracellular space that acts as a paracrine signaling molecule, facilitating cell-to-cell communication through the propagation of calcium signals ^107–109^. Thus, the depletion of macrophages within the pituitary gland could potentially disrupt calcium homeostasis by reducing the availability of extracellular ATP. The outstanding challenge now for the future is to define the intercellular communication between macrophages and different hormone-producing cells in the delicate 3D environment of the pituitary gland.

In summary, we present a transcriptomic characterization of pitMØ, revealing diverse features. At a steady state, the pituitary gland hosts macrophages of diverse characteristics in different spatial localizations, ml-MACs expressing microglia-like genes in the posterior and other pitMØ in the anterior pituitary gland interacting closely with hormone-secreting cells. All pitMØ originate from early yolk sac progenitors and self-renew without input from bone marrow-derived monocytes. Our findings strongly suggest the importance of macrophages in the pituitary gland and highlight their roles in hormone production and intercellular communication. However, further research into the specific mechanisms by which macrophages exert these effects will provide valuable insights into pituitary physiology and may uncover potential therapeutic targets for hormone-related disorders.

## Methods

### Mice

Multiple genetic mouse models were used in this study: *Ccr2^−/−^*(stock 004999), *R26^EFGP^* (stock 006148), *R26^tdTomato^* (stock 007914), and *Cx3cr1^CreERT^*^2^ (stock 020940) were purchased from Jackson Laboratories. *CCR2^CreERT^*^2^ and *IL34^lac^* mice were kindly provided by Prof. Dr. Burkhard Becker, *Ms4a3^CreERT^*^2^ mice were kindly provided by Prof. Florent Ginhoux, and *Rb1^tm1Tyj^*mice were kindly provided by Prof. Jorma Toppari. Wild-type (WT) mice, C57BL/6N, were acquired from Janvier labs. Experiments with 1-week and older were performed only with male mice. In embryonic and NB timepoints the whole litter was pooled for pituitary gland samples. All experimental mice were kept in specific pathogen-free conditions and under controlled environmental conditions at a temperature of 21 °C ±3 °C, humidity 55 % ±15 %, and 12/12 h light cycle at the Central Animal Laboratory of the University of Turku (Turku, Finland). Mice had food and water *ad libitum*. Animal experiments were conducted under the revision and approval of the Regional Animal Experiment Board in Finland, according to the 3R principle, and under Animal license numbers 6211/04.10.07/2017 and 14685/2020. All experiments were regulated according to the Finnish Act on Animal Experimentation (497/2013). Embryonic development was estimated considering the day of a vaginal plug as embryonic day 0.5 (E0.5).

### Single cell isolation

The pregnant females were euthanized by carbon dioxide (CO2) asphyxiation followed by cervical dislocation. Embryos were dissected from the uterus and euthanized through decapitation. Newborn and 7-day-old pups were euthanized by decapitation. From 2 weeks of age onwards, euthanasia was carried out using CO2 asphyxiation followed by cervical dislocation or cardiac puncture. For the ScRNA sequencing, 3- and 8-week-old male mice were sacrificed with CO_2_, the chest cavity was opened by two parasternal incisions, and the perfusion was started immediately. The perfusion was done through the left ventricle using a peristaltic pump (Harvard peristaltic Pump P-230, Harvard apparatus) with a constant pressure of 5 ml/min for 3 min with cold PBS. The right atrium was opened for the outflow. The whole pituitaries were collected. For ScRNA sequencing and flow cytometry, whole pituitaries or separate posterior and anterior pituitaries were mechanically lysed by suspending and digested with 50 µg/ml DNase 1 (Roche) and 1 mg/ml Collagenase D (Roche) in Hank’s Buffered Saline (HBSS; Sigma-Aldrich) at 37 °C for 30-45 minutes. The cell suspension was filtered through silk (pore size 77 µm), pelleted (1006G, 1.5 minutes), and washed with 500 µl of HBSS to remove enzymes. Finally, cells were eluted to Phosphate Buffered Saline (PBS; Sigma-Aldrich) before labeling for cell sorting or analyzing flow cytometry.

Blood from mouse embryos to 1 week of age mice was collected by bleeding the body to heparin-PBS (50 µl of 100 IU/ml heparin in 500 µl of PBS). From 2 weeks of age onwards, blood was drawn by cardiac puncture into heparinized syringes, and erythrocytes were lysed from the blood as described ^110^.

Brain, liver, and lungs were carefully dissected, minced with scissors, and digested with 50 µg/ml Dnase 1 and 1mg/ml Collagenase D in HBSS at 37 °C (20 min for brain and 60 min for liver and lung) and then passed through a 70-μm cell strainer. The brain cells were then resuspended in isotonic Percoll (Sigma-Aldrich), and the microglia were isolated as described ^52^. Finally, cells were eluted to PBS before labeling for flow cytometry.

### ScRNA sequencing and data analysis

Single cell suspensions derived from WT pituitary glands at NB, 1, 3, and 8 weeks of age were stained with Fixable Viability Dye eFluor 780 (eBioscience) to label the dead cells (Pooled samples of NB n=31, 1 week n=15, 3 weeks n=25 and 8 week n=21 animals). Unspecific binding to low-affinity Fc-receptors was blocked by incubating the cells with unconjugated CD16/32 antibody (BioXCell, clone 2.4G2). Cells were subsequently stained for 30 min at 4 °C with antibodies diluted to the FACS buffer. The cells were sorted with Sony SH800 Cell Sorter (Sony Biotechnology Inc.). Single cells were gated with the FSC-H versus FCS-W plot and the SSC-H versus SSC-W plot to avoid doublets and dead cells were excluded. Live single CD45^+^ cells were sorted into the RPMI medium containing 2 % FCS.

Freshly sorted CD45^+^ were processed immediately according to 10X Genomics guidelines (CG000126_Guidelines for Optimal Sample Prep Flow Chart RevA). Single cell RNA-sequencing libraries were prepared according to the manufacturer’s instructions (CG00052 RevB) using Chromium Single Cell 3′ Library and Gel Bead Kit v2 (10X Genomics, 120237) and Chromium Single Cell A Chip Kit (10X Genomics). Prepared libraries were sequenced using the Illumina Novaseq6000 Sequencing System (RRID: SCR_016387). Library preparation was done in the Single Cell Omics core, and sequencing was done in the Finnish Functional Genomics Center at Turku Bioscience. The library preparation was executed at the Single Cell Omics core, and the sequencing was conducted at the Finnish Functional Genomics Center, both housed within Turku Bioscience Center, Finland.

The *Cell Ranger* (ver. 7.1.0) “count” function was used to align samples to the *mm10-2020-A* reference genome, quantify reads, and filter reads and barcodes. Spliced/unspliced counts for RNA velocity estimation were calculated with the Python implementation of *Velocyto* (ver. 0.17.17, SCR_018167 ^111^). Data analysis and displays were generated using the *scanpy* (ver. 1.9.8, SCR_018139 ^112^), *scVelo* (ver. 0.3.1, SCR_018168 ^113^), *CellRank* (ver. dev2.0.2, SCR_022827 ^114^), and Python (ver. 3.10, SCR_008394) toolboxes.

For quality control, low-quality cells were filtered out in each sample by the number of expressed unique genes and percentage of mitochondrial genes (max.: 8 %). Genes with less than 20 shared spliced/unspliced counts and genes expressed by fewer than 3 cells were removed. Counts per cell were normalized as total counts over all genes (*target sum: 10^4*) and were log1p transformed. Principal components (PCA) were calculated and used for sample integration with harmony (ver. 0.0.6 ^115^). For the complete dataset, the neighborhood graph (nearest neighbors: 50), Uniform Manifold Approximation and Projection (UMAP; SCR_018217 ^116^; min. distance: 0.5, neg. edge sample rate: 8), and Leiden clustering (ver. 0.9.1 ^117^; resolution: 1.8) were computed with the indicated settings.

Clusters containing monocytes and macrophages were selected by marker gene expression, and neighborhood graph (*nearest neighbors: 50*), UMAP (*min. distance: 0.7, neg. edge sample rate: 12*), and Leiden clustering (*resolution: 0.75*) were re-computed. Transcriptional dynamics were re-constructed using scVelo’s generalized dynamical model, and CellRank’s CytoTRACE Kernel (*threshold scheme: soft*) was used to derive a pseudotime ordering of cells based on cellular plasticity estimation. Differential expression genes were computed with sci-tools’s (ver. 1.0.4) single cell Variational Inference (scVI; *layer: counts, categorical covariates: batch, continuous covariate: pct_counts_mt*) model ^118^ and the *differential_expression* function (*mode: “change’, delta: 0.25, fdr_target: 0.05*; ^119^).

### In utero and adult tamoxifen administration

*Cx3cr1^CreERT^*^2^ and *CCR2^CreERT^*^2^ mice were crossed with *R26^EYFP^*or *R26^tdTomato^* reporter mice to study embryonic-derived macrophages. To study bone marrow-derived macrophages, *Ms4a3^CreERT^*^2^ mice were crossed with the *R26^tdTomato^* reporter strain. For *in utero* tamoxifen induction of CRE activity, a single dose of tamoxifen (1.5 mg/dam, mended with 0.75 mg progesterone; Sigma–Aldrich) was administered i.p. to pregnant females on indicated pregnancy days. For adult tamoxifen induction, *Cx3cr1^CreERT^*^2^*; R26^EYFP^*or *Ms4a3^CreERT^*^2^*; Rosa^tdTomato^* mice were administered i.p. 5 consecutive days of tamoxifen (75 mg tamoxifen/kg). This induction scheme leads to selective labeling of CX3CR1 positive macrophages and monocytes or bone marrow-derived monocytes.

### Flow cytometry

Cells were stained with Fixable Viability Dye eFluor 780 (eBioscience) to label the dead cells. Unspecific binding to low-affinity Fc-receptors was blocked by incubating the cells with unconjugated CD16/32 antibody (BioXCell, clone 2.4G2). Cells were subsequently stained for 30 min at 4 °C with antibodies diluted to the FACS buffer, washed, and fixed with 1 % formaldehyde in PBS. Samples were acquired with LSRFortessa flow cytometer with FACSDiVaTM version 9.0 software (Becton Dickinson), and data were analyzed using FlowJo™ v10.8 Software (BD Life Sciences)

### Immunofluorescence staining and confocal microscopy

For whole mount stainings, pituitary glands were collected from male mice at 17.5, new born, 1 week, 2 weeks, 5 weeks, and 8 weeks of age. Pituitaries were dissected attached to the skull to prevent the gland from breaking. Samples were prepared for whole mount staining by fixing them in 2 % paraformaldehyde (PFA, Santa Cruz Biotechnology) in PBS for 20 minutes on ice. Samples were washed in PBS (3x 10 min) shaking. The pituitary was dissected from the skull between the second and third wash. Whole mount samples were dehydrated in a graded methanol series and stored at -20 °C. Samples were rehydrated through ascending series of methanol and finally placed into PBS. In brief, samples were blocked [1 % normal goat serum (NGS) + 0.5 % fetal calf serum (FCS) + 1 % bovine serum albumin (BSA) + 0.4 % Triton-X 100 in PBS] and then sequentially incubated with primary unconjugated antibodies, secondary antibodies and directly conjugated antibodies diluted in blocking solution. Finally, the samples were dehydrated with increasing methanol series and subsequently optically cleared in glass-bottom microwell dishes first with 50 % benzyl alcohol (Honeywell) 1:2 benzyl benzoate (BABB; Sigma–Aldrich) in methanol and then with 100 % BABB, which was also used as a mounting medium.

For immunofluorescence stainings, pituitaries were embedded in OCT medium (Tissue-Tek), freezed on dry ice, and stored at −80 °C prior to cryosectioning. Pituitaries were cut into 7-µm-thick sections and fixed with ice-cold acetone for 5 minutes. Before staining, samples were post-fixed in 2 % PFA for 15 min at room temperature (RT) and rinsed in PBS. Samples were blocked for 1 hour at RT in a humidity chamber in a blocking buffer (1 % FCS, 1 % NMS, and 5 % NGS in 0.3 % Triton X-PBS). Primary antibodies used for LHβ (1:1000, anti-mouse rabbit, Parlow), TSH (1:1000, Parlow), or PRL (1:1000, Parlow) were diluted in blocking solution and applied to sections overnight at 4 °C. Sections were then washed 2 times 3 minutes in 0.1 % TritonX-PBS. After washes, a fluorescent secondary antibody was added for 60 minutes at RT, followed by washes in 0.1 % TritonX-PBS for 3 times 5 minutes and a rinse in PBS. Sections were counterstained with DAPI (1:5000 in PBS, 25 µl per section) and incubated for 10 minutes in a humidity chamber at RT. Sections were rinsed in PBS and MQ water. Finally, slides were mounted with the Mounting Prolong Gold antifade without DAPI (ThermoFisher Scientific).

Imaging was performed using an LSM880 confocal microscope (Zeiss) with an Axio Observer.Z1 microscope using ZEN 2.3 SP1 black edition acquisition software at RT. The objectives used were a Zeiss Plan-Apochromat 20x/0.8 without immersion, Zeiss LD LCI Plan-Apochromat 25x/0.8 with glycerol immersion, and Zeiss LD LCI Plan-Apochromat 40x/1.2 with water immersion. Further image processing was performed with ImageJ software (National Institute of Health).

### X-gal staining

For X-gal staining, 5-week-old male mice were sacrificed with CO_2_, the chest cavity was opened by two parasternal incisions, and the perfusion was started immediately. Mice were perfused first with 10 ml of PBS and then with 10 ml of 2% PFA in PBS. The right atrium was opened for the outflow. After collection, the pituitaries were fixed in 2 % formaldehyde and 0.2 % in PBS solution for 15 minutes. After fixation, the pituitary glands were embedded into OCT blocks and frozen at -80 °C. Pituitaries were cut into 6 µm sections with cryotome and fixed in acetone for 5 minutes. Staining was done with a β-gal staining kit (Invitrogen), according to the manufacturer’s instructions, with the X-gal dissolved in DMSO (Sigma-Aldrich). Exceptions were that slides were washed 3 times with PBS after fixation, and after staining, washing was done with 1 x PBS and water. Slides were dehydrated with 96 % ethanol for 30 seconds. Counterstaining was done with 0.1 % eosin solution (180072, Reagena) for 3 minutes, after which slides were dehydrated in ethanol and xylene series. Mounting was done with DPX mountant for histology (Sigma-Aldrich). Slides were imaged with Pannoramic P1000 (3DHISTECH) and analyzed with Pannoramic Viewer (Version 1.15.4, 3DHISTECH).

### LPS treatment

A single injection of Lipopolysaccharides (LPS) from *Escherichia coli* (O55:B5; Merck; 20 µg) was given i.p. for adult 8-10 weeks old mice. Control mice were injected with the same volume of PBS. The pituitary glands were analyzed by flow cytometry 24 hours later. For treatment control, blood was collected from the treated mice by cardiac puncture, and serum was separated by centrifugation. IL-6 was analyzed from blood serums by sandwich enzyme-linked immunosorbent assay (ELISA; Thermo Fisher Scientific) according to the manufacturer’s instructions.

### Antibody treatment experiments

To specifically target yolk sac-derived macrophages in the offspring, pregnant C57BL/6N females at embryonic day 6.5 (E6.5) were treated with a single intraperitoneal (i.p.) injection of a neutralizing antibody against CSF1R (AFS98; Bio X Cell) or received rat IgG2a control antibody (clone 2A3; Bio X Cell). Each injection consisted of 3 mg of the respective antibody dissolved in sterile PBS. The pituitary glands, testis, and brains were collected at E17.5 for flow cytometric analyses.

A single injection of function-blocking CSF1 antibody (5A1; Bio X Cell; 150 µg), preventing the binding to CSF1R, or a neutralizing antibody to CSF1R (150 µg) was given i.p. for newborn mice. Control mice were injected with control IgG (HRPN; Bio X Cell; 150 µg) or rat IgG2a control antibody (2A3; Bio X Cell; 150 µg). The pituitary glands were analyzed by flow cytometry 48 hours later. The brain was used as treatment control tissue.

To address the functional importance of macrophages for normal pituitary gland function, we ablated the pre-existing tissue-resident macrophages with CSF1R and gave the first injection during embryo development at E6.5. In a long depletion experiment, CSF1R was additionally given on postnatal day (P) 2, P14, and 4 weeks of age (4 times in total). The pituitary glands and brains were collected at 5 weeks of age.

### Histological analyses

Pituitary glands were dissected attached to the skull and fixed with 10 % Formalin (Sigma-Aldrich) at RT overnight. After fixing, pituitary glands were dehydrated through graded ethanol (EtOH) series at RT. 5 µm thick sections were cut and stained with Mayers hematoxylin (Sigma-Aldrich) and eosin (Reagena). Slides were mounted with DPX mountant for histology (Sigma-Aldrich), imaged with Pannoramic P1000 (3DHISTECH), and analyzed with Pannoramic Viewer (Version 1.15.4, 3DHISTECH).

### Bulk RNA sequencing and data analysis

To determine the gene expression signature of macrophage-depleted pituitaries, we performed bulk RNA-seq from the total RNA of the pituitary glands collected after long CSF1R antibody treatment (see protocol above). RNA was isolated with RNeasy Plus Micro Kit (Qiagen) according to the manufacturer’s protocol and analyzed on Advanced Analytical Fragment Analyzer for quality assessment. The sequencing run was performed using Illumina NovaSeq 6000 SP v1.5 (650-800 M reads/run, 2 lanes; Illumina) in the Finnish Functional Genome Centre (Turku Bioscience Center, Finland). The quality of the sequenced reads was investigated. The reads were aligned to the mouse reference genome (GRCm38 release 94) using HISAT2 v2.0.5 ^120^. The novel gene assembly was predicted using StringTie v1.3.3b ^121^ using the mapping information from all the samples and the novel gene annotation was performed using Pfam, Swissprot, Go, and KEGG. The number of uniquely mapped reads associated with each gene was counted using the featureCounts v1.5.0-p3 ^122^ tool using Ensembl annotations. The downstream analysis of gene counts was performed using R v4.1.2 ^123^ and corresponding Bioconductor v3.14 ^124^. The lowly expressed genes were filtered by using filterByExpr in edgeR package ^125^. To tackle the issue of batch effects, we corrected the gene counts using ComBat_seq function ^126^ in the SVA R package ^127^ which is specially designed for RNA-seq data. It corrects for the batch effect by fitting a negative binomial regression model. The batch-corrected counts were normalized for library size and statistically tested using the DESeq2 package ^128^. The genes were considered differentially expressed if their adjusted p-value, estimated using the Benjamini-Hochberg test was equal to or lower than 0.1 and absolute fold change was greater than or equal to 2 between the group comparison between IgG2a and CSF1R. Gene set enrichment analyses (GSEA) were performed using the GSEA v 4.3.3 (Broad Institute, Cambridge, MA) with the M5; GO: BP (containing 7796 gene sets) and M2; CP; REACTOME (containing 1261 gene sets) collections from the molecular signature database (https://www.gsea-msigdb.org/gsea/msigdb/index.jsp). Analysis was performed in January 2024, with a significant P value <0.05 and a false discovery rate (FDR) <0.25 ^129–131^. In addition, ingenuity pathway analysis (IPA) was performed using IPA software (Qiagen, CA, USA). IPA analysis was performed using DE genes with a significant P value <0.05 and fold change >1.6.

### Hormone measurements

For the intrapituitary hormone measurements, pituitaries were homogenized in 500 µl phosphate-buffered saline (PBS). Concentrations of ACTH, LH, FSH, GH, TSH, and PRL were analyzed in a single run on a Luminex 200 platform using a MILLIPLEX MAP Mouse Pituitary Magnetic Bead Panel (Millipore) according to the manufacturer’s instructions. Intra- and inter-assay coefficients of variation (CV) were lower than 12.4 %. The lowest detection limits for intrapituitary ACTH, LH, FSH, GH, TSH, and PRL measurements were 1.7, 4.9, 9.5, 1.7, 1.9, and 46.2 pg/ml, respectively.

### mRNA expression by quantitative real-time PCR

Tissues were snap-frozen with liquid nitrogen, and total RNA was extracted. DNAse treatment was carried out with RNeasy Plus Micro/Mini Kit (Qiagen) following the manufacturer’s instructions. RNA samples were dissolved in water, and after the quality check, RNA was used for cDNA synthesis using a SensiFast cDNA synthesis kit (Bioline). qPCR reactions were performed with TaqMan™ Universal Master Mix II (Applied Biosystems™). Gene expression was quantified by QuantStudio™ 3 or 12K Flex Real-Time PCR Systems (Thermo Fisher Scientific). The mRNA expression of *Gnrhr* was normalized to the expression of *Actb*, and the expression of mRNA was quantified using the PfaffI method ^132^. The sequences of the primers used for *Gnrhr* were 5′-TGCTCGGCCATCAACAACA-3′ and 5′-GGCAGTAGAGAGTAGGAAAAGGA-3′, and for *Actb*, 5′-CGTGGGCCGCCCTAGGCACCA-3′ and 5′-TTGGCCTTAGGGTTCAGGGGG-3′. The expression of *Gnrhr* was analyzed in the pituitary glands of CSF1R- and IgG control-treated mice (n = 4).

### Statistical analyses

Adult mice were assigned to experimental groups without employing specific randomization methods, as comparisons involved mice with varying genetic backgrounds. The researchers conducted the experimental procedure while blinded to the genotype of the embryos. Statistical analyses were conducted using GraphPad Prism software version v10.1.2 (GraphPad Software Inc). All data are presented as mean values ± SEM. Comparisons between the time points, genotypes, or treatment groups were made using the nonparametric two-tailed Mann-Whitney test, parametric two-tailed t-test, and one-way ANOVA test with Bonferroni post-hoc test. P-values lower than 0.05 were considered statistically significant. The value of each n can be found in the Figure legends.

## Supplemental information

**Figure S1.**
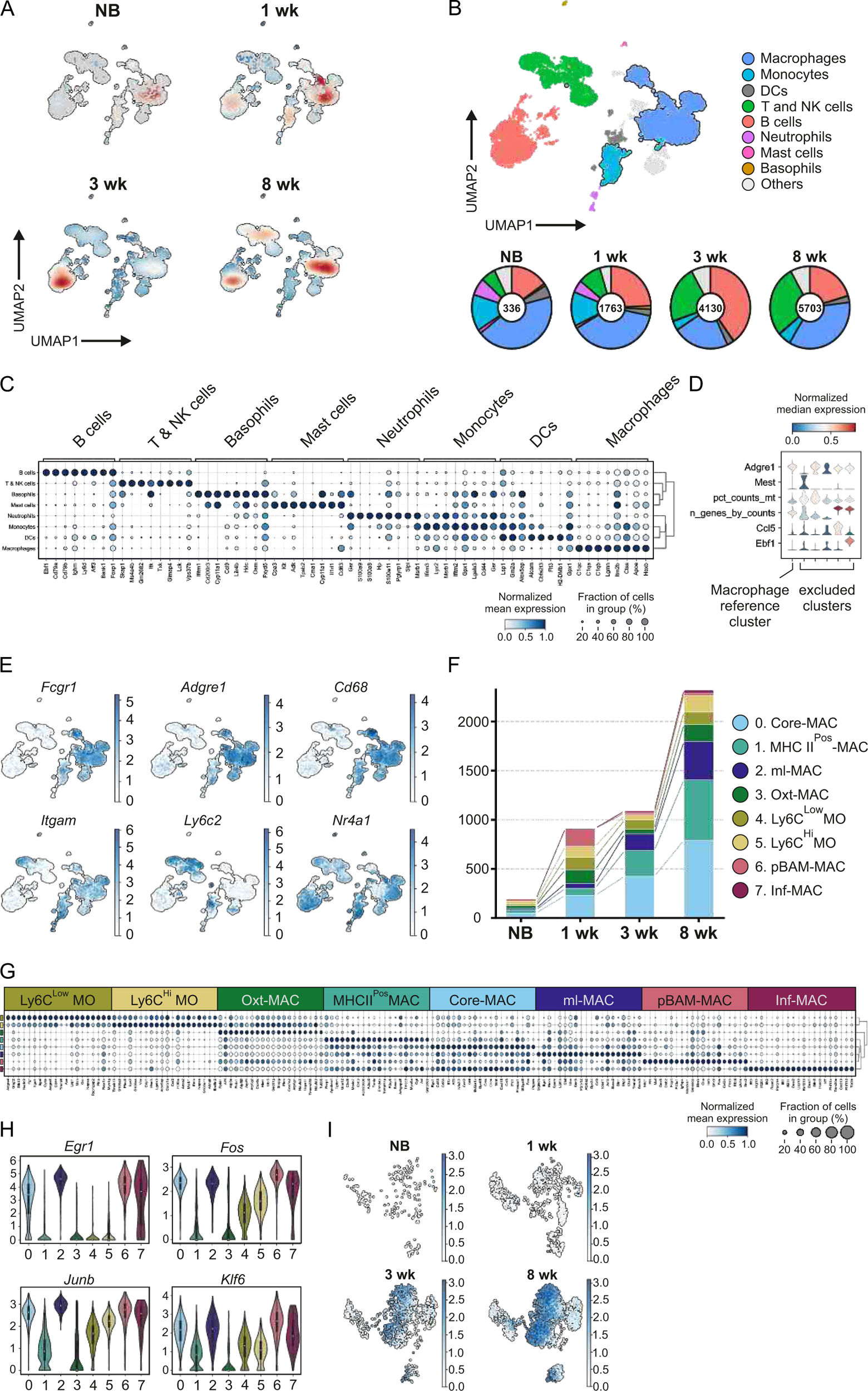
Heterogeneity of leukocytes (CD45^Pos^ cells), pitMØ and monocytes in mouse pituitary gland, related to figure 1. (A) UMAP plots highlighting the position of cells on the UMAP plot across time points. Color intensity represents cell embedding density from low (blue) to high (red). (B) UMAP plot of mouse pituitary gland leukocytes (CD45^Pos^ cells) colored by the cluster. Donut charts present the cell frequency of different leukocyte populations per sampling time point in the developing pituitary gland. (C) Dotplot of the top-ranked genes in each leukocyte population. The color code indicates the expression level, from low (white) to high (dark blue). (D) Violin plots of gene expression and sample properties used for cluster selection of reference macrophage and excluded *Leiden* clusters colored by row normalised median feature values. (E) The expression of selected genes on the UMAP plot of mouse pituitary gland leukocytes. The color code indicates the expression level from low (white) to high (dark blue). (F) Stacked bar charts present the cell amount of pitMØ and monocytes at different time points (NB, 1 wk, 3 wk, and 8 wk). (G) Dotplot of single-cell expression of the top 20 ranked genes in pitMØ and monocytes. The color code indicates the expression level from low (white) to high (dark blue). (H) Violin plots showing expression of selected immediate early genes in different pitMØ and monocyte clusters. (I) The expression level of *H2-ab1* (gene coding MHC II) on the UMAP plot of pitMØ and monocytes of the mouse pituitary gland. The color code indicates the expression level from low (white) to high (dark blue).

**Figure S2.**
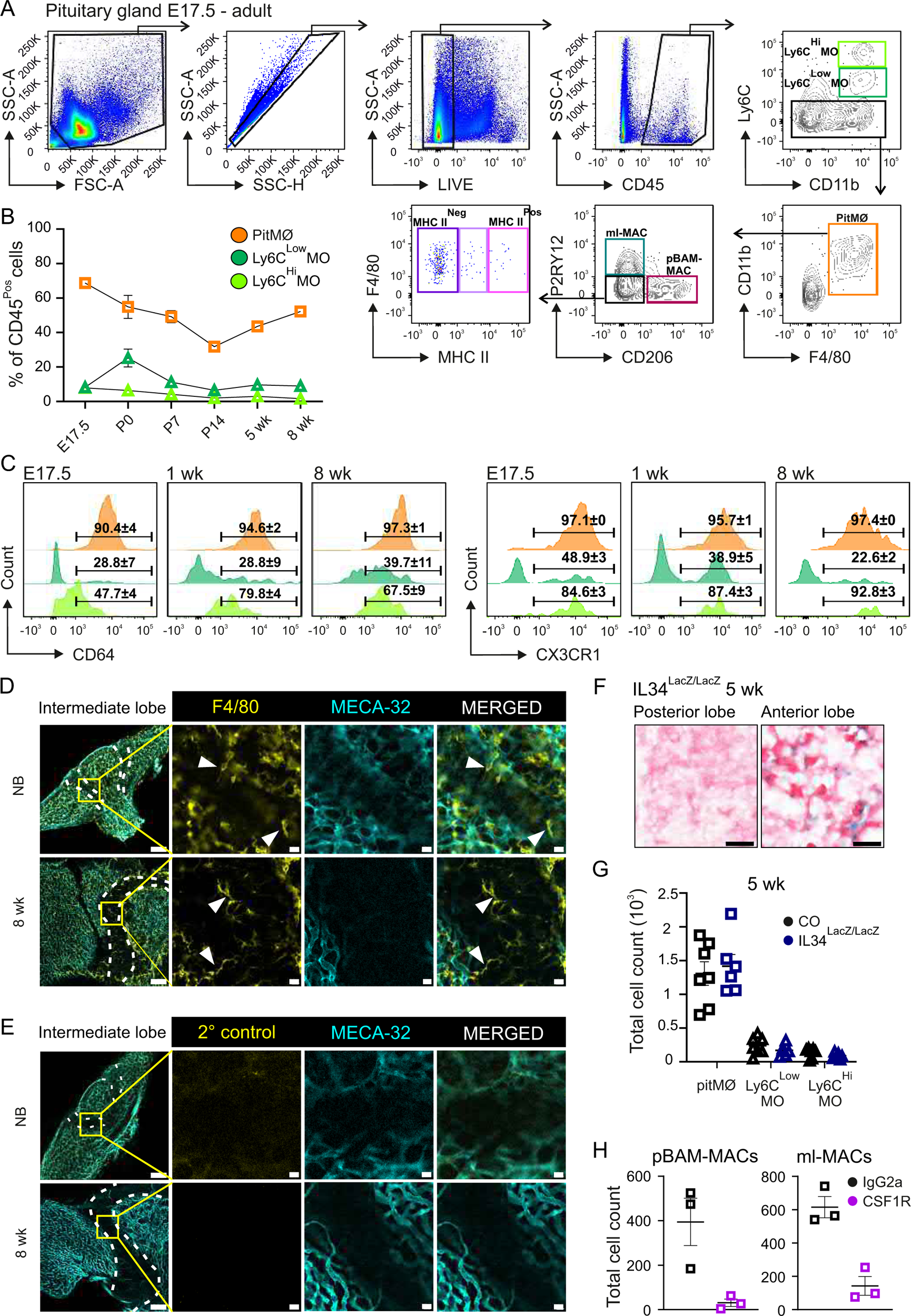
Development modifies the phenotype of CSF1R-dependent pitMØ, related to figure 2. (A) Gating strategy for pitMØ and monocytes in all timepoints. (B) Frequency of pitMØ, Ly6C^Low^MO and Ly6C^Hi^MO cells in CD45^Pos^ cells of the WT mouse pituitary gland at indicated timepoints. The quantitative data are shown as mean ± SEM. Each dot represents data from mice pooled together in E17.5 (n=8-12 samples/pool), NB (n=7-11 samples/pool), 1 wk (n=4-6 samples/pool), 2 wk (n=2-5 samples/pool), and from individual mice at 5 wk (n=15 mice) and 8 wk (n=13 mice). Data are from 6-8 individual experiments. (C) Representative histograms of CD64 and CX3CR1 expression in pitMØ, Ly6C^Low^MO and Ly6C^Hi^MO populations at indicated timepoints in WT mice. Each dot represents data from mice pooled together in E17.5 (n=9-12 samples/pool), 1wk (n=4-6 samples/pool), and from indivicual mice at 8 wk (n = 4-9). Data are from 2-4 individual experiments. (D) Pituitary gland whole mount immunofluorescence with F4/80 and endothelial marker, MECA-32, from WT pituitary glands of NB and 8-week-old mice. Inserts are higher magnifications from the boxed intermediate lobe areas. Arrow heads indicate F4/80^pos^ macrophages locating in the edge of intermediate lobe. For overview images, scales are 100 µm, and for inserts, 10 µm. (E) Pituitary gland whole mount immunofluorescence with secondary antibody, anti-rat IgG goat A546, and endothelial marker, MECA-32, from WT pituitary glands of NB and 8-week-old mice. Inserts are higher magnifications from the boxed intermediate lobe areas. For overview images, scales are 100 µm, and for inserts, 10 µm. (F) IL34 expression by X-gal staining in the anterior and posterior pituitary gland. Scale 20 µm. Shown are representative images from 3 independent experiments. (G) The cell amount of pitMØ, Ly6C^Low^MO and Ly6C^Hi^MO cells in *Il-34* deficient mouse pituitary gland at 5 weeks of age. The quantitative data are shown as mean ± SEM. Each dot represents data from one mouse (n=6-7). Data are from 3 individual experiments. (H) The cell amount of CD206^Pos^ and P2RY12^Pos^ cells at postnatal day (P) 2 after treating NB WT mice with CSF1R antibody or control IgG at NB. The quantitative data are shown as mean ± SEM. Each dot represents data from mice pooled together (n=7-11 samples/pool). Data are from 5 individual experiments.

**Figure S3.**
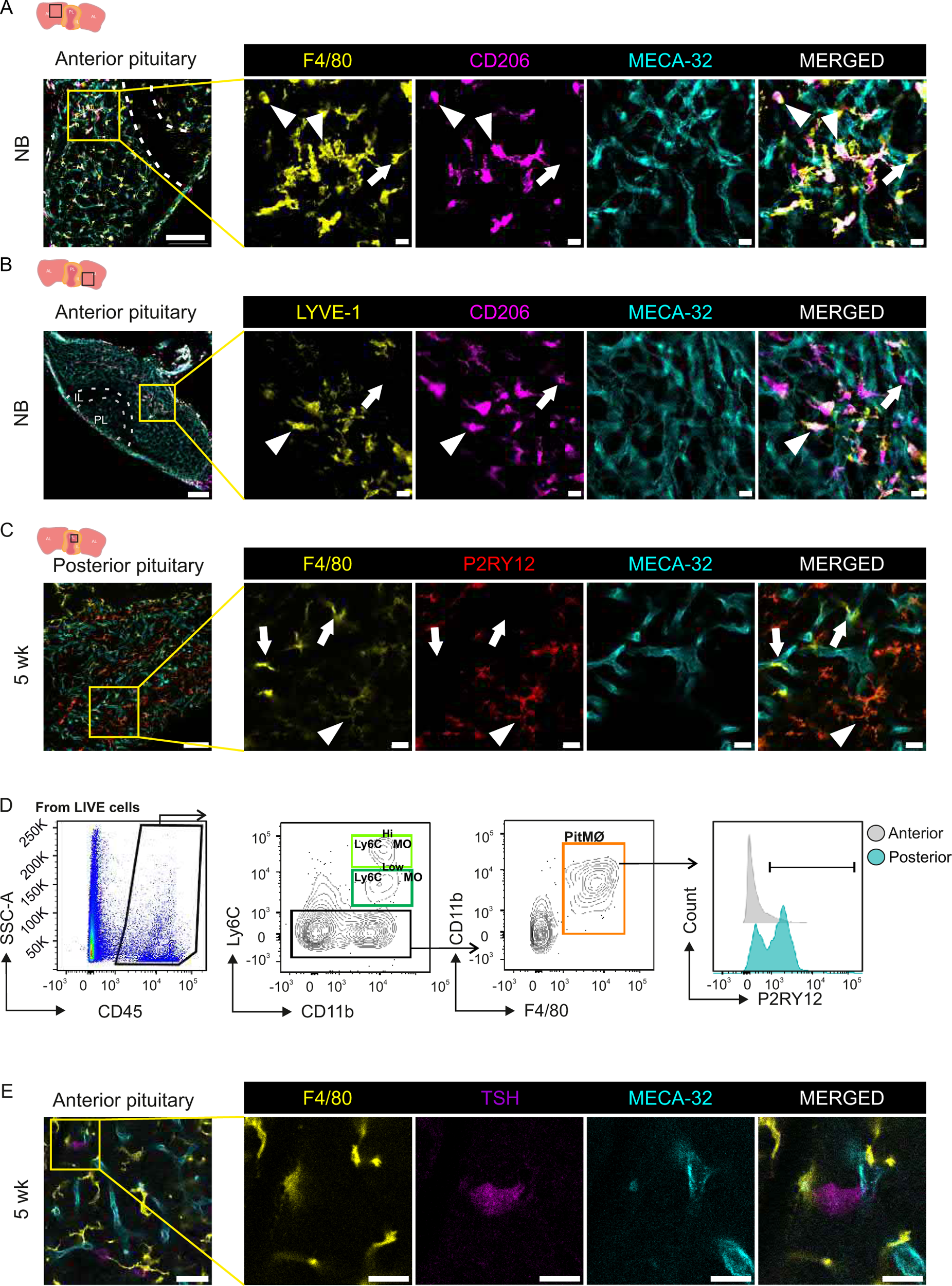
Localization of CD206, LYVE-1 and P2RY12 in the pituitary gland, related to figure 3. (A) Pituitary gland whole mount immunofluorescence with F4/80, CD206 and endothelial marker, MECA-32, from WT pituitary glands of NB mice. Inserts are higher magnifications from the boxed anterior lobe area. Arrows indicate F4/80^Pos^CD206^Neg^ cells and arrowheads indicate F4/80^Pos^CD206^Pos^ cells. For overview image, scales is 100 µm, and for inserts, 10 µm. Shown are representative images from 3 independent experiments. (B) Pituitary gland whole mount immunofluorescence with LYVE-1, CD206 and MECA-32 from WT pituitary glands of NB. Inserts are higher magnifications from the boxed anterior lobe area. Arrows indicate LYVE-1^Neg^CD206^Pos^ cells and arrowheads indicate LYVE-1^Pos^CD206^Pos^ cells. For overview image, scales is 100 µm, and for inserts, 10 µm. Shown are representative images from 3 independent experiments. (C) Pituitary gland whole mount immunofluorescence with F4/80, P2RY12 and MECA-32 from WT pituitary gland of 5 weeks of age. Inserts are higher magnifications from the boxed posterior lobe area. Arrows indicate F4/80^Pos^P2RY12^Neg^ cells and arrowheads indicate F4/80^Pos^P2RY12^Pos^ cells. For overview image, scales is 50 µm, and for inserts, 10 µm. Shown are representative images from 3 independent experiments. (D) Gating strategy and representative histograms of P2RY12 expression in anterior and posterior lobes from the pituitary gland of 10-week-old mice. The quantitative data are shown as mean ± SEM. Each dot represents data from one mouse (n=3-8). Data are from 2 individual experiments. (E) Pituitary gland whole mount immunofluorescence with F4/80, thyroid-stimulating hormone (TSH) and endothelial marker, MECA-32, from WT pituitary gland of 8 weeks of age. Inserts are higher magnifications from the boxed anterior areas. For overview image, the scale is 100 µm, and for inserts, 10 µm. Shown are representative images from 3 independent experiments.

**Figure S4.**
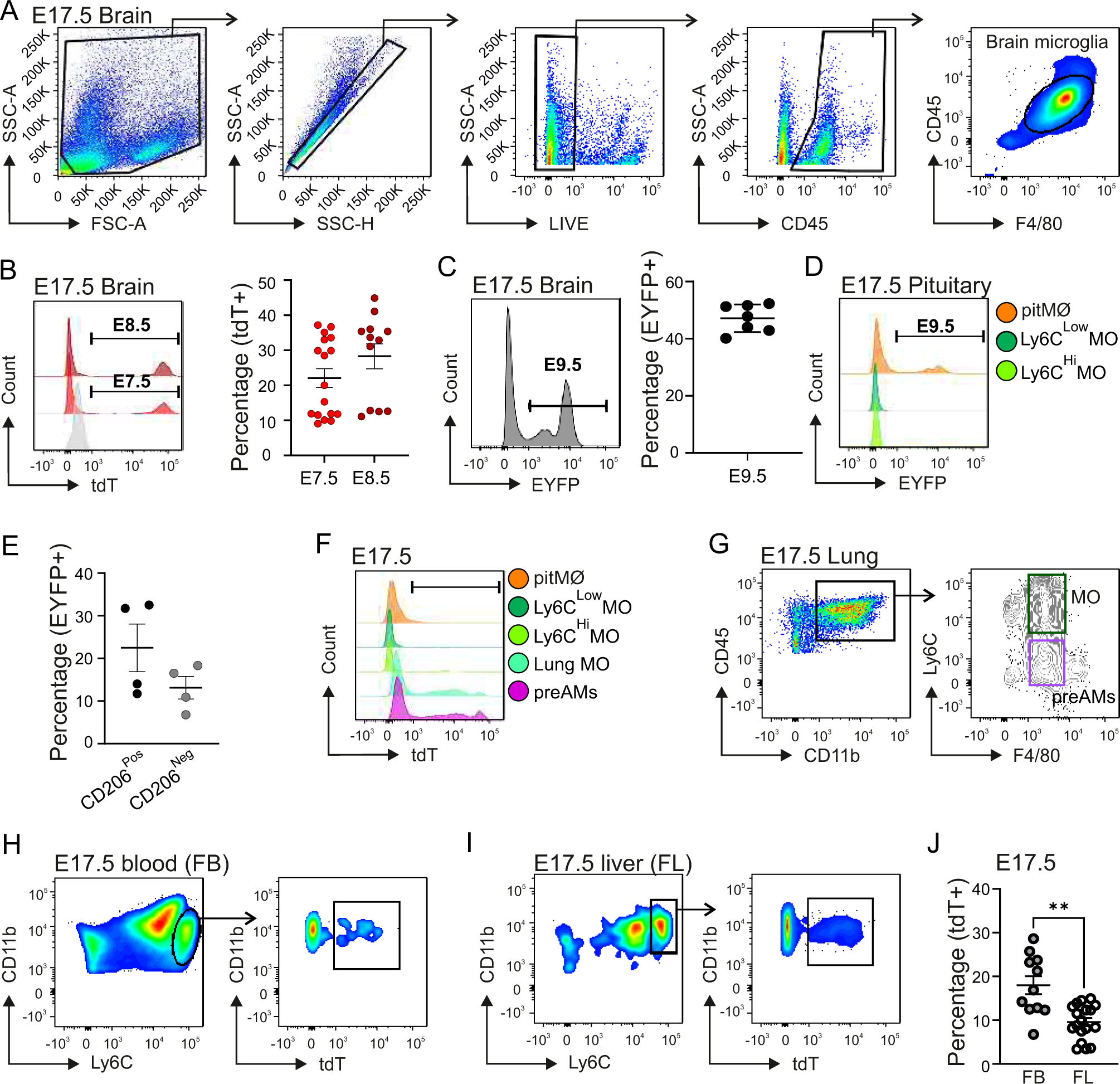
Pituitary gland macrophages originate from early yolk sac progenitors, related to figure 4. (A) Gating strategy for E17.5 brain microglia. (B) Representative histograms and frequencies of tdTomato expression in microglia of E7.5 and E8.5 tamoxifen labelled Cx3cr1^CreERT2^R26^tdTomato^ brains at E17.5. The quantitative data are shown as mean ± SEM. 3 experiments, each symbol represents one animal (n=12-17). (C) Representative histogram and frequency of EYFP^Pos^ cells in brain microglia of E9.5 tamoxifen labelled Cx3cr1^CreERT2^R26^EYFP^ brains at E17.5. The quantitative data are shown as mean ± SEM. Each dot represents data from one mouse. Each dot represents data from mice pooled together (n=2-3 samples/pool). Data are from 2 individual experiments. (D) Representative histogram of EYFP expression in pitMØ, Ly6C^Low^MO and Ly6C^Hi^MO populations of E9.5 tamoxifen labelled Cx3cr1^CreERT2^R26^EYFP^ pituitary glands at E17.5. Each dot represents data from mice pooled together (n=9-11 samples/pool). Data are from 4 individual experiments. (E) Frequency of EYFP^Pos^ expression in CD206^Pos^ (pBAM-MACs) and CD206^Neg^ cells in pituitary glands at E17.5. Each dot represents data from mice pooled together (n=9-11 samples/pool). Data are from 4 individual experiments. (F) Representative histogram of tdTomato expression in pitMØ, Ly6C^Low^MO, Ly6C^Hi^MO, Lung MO and preAMs populations of E14.5 ans E16.5 tamoxifen labelled Ccr2^CreERT2^R26^EYFP^ at E17.5. Each dot represents data from mice pooled together (pitMØ, Ly6C^Low^MO, Ly6C^Hi^MO, n=8-10 samples/pool) or one animal (Lung MO, preAMs, n=19). Data are from 4 individual experiments. (G) Gating strategy for E17.5 lung monocytes and pre-alveolar macrophages (preAMs). (H) Gating strategy for E17.5 tdTomato^Pos^ cells from blood monocytes (FB). (I) Gating strategy for E17.5 tdTomato^Pos^ cells from liver monocytes (FL). (J) Frequency of tdTomato^Pos^ cells in blood (FB) and liver (FL) monocytes. Each dot represents data from one mouse (n=11-18). Data are from 4 individual experiments.

**Figure S5.**
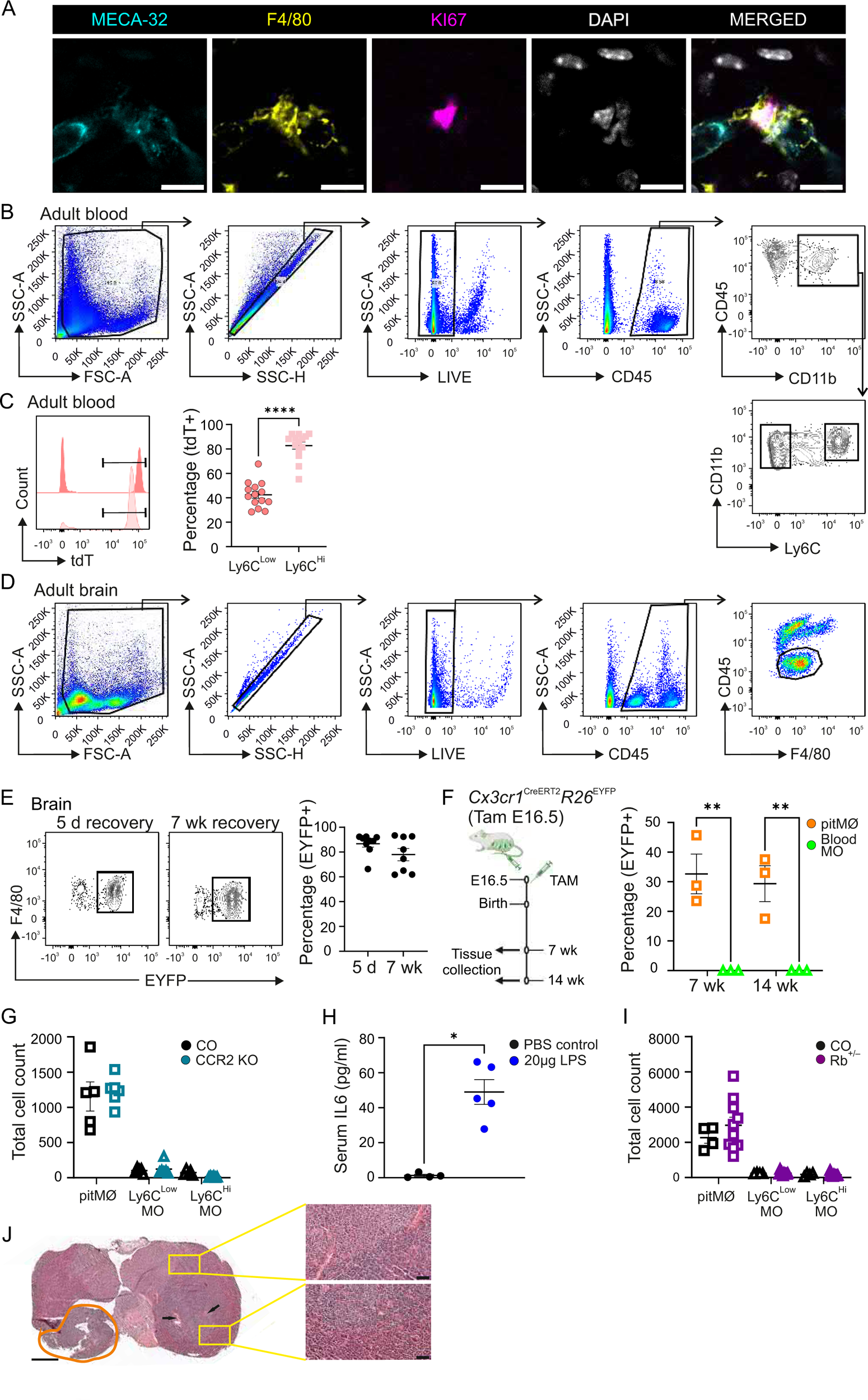
Early yolk sac-derived pituitary gland macrophages self-maintain until adulthood by proliferation, related to figure 5. (A) Pituitary gland whole mount immunofluorescence with F4/80, proliferation marker, KI67, endothelial marker, MECA-32, and nuclear marker DAPI from WT pituitary glands of 1-week-old mice. Scales are 5 µm. Shown are representative images from 3 independent experiments. (B) Gating strategy for adult blood monocytes (Ly6C^Hi^ and Ly6C^Low^). (C) Representative histograms and frequency of tdTomato^Pos^ cells in blood Ly6C^Low^ and Ly6C^Hi^ monocytes of Ms4a3^CreERT2^R26^tdTomato^ mice at 5 weeks of age (4 day recovery). Tamoxifen induction for five consecutive days. The quantitative data are shown as mean ± SEM. Each dot represents data from one mouse (n=14). Data are from 4 individual experiments. (D) Gating strategy for adult brain microglia. (E) Representative flow cytometry plots and frequency of EYFP^Pos^ cells in brain microglia in the Cx3cr1^CreERT2^R26^EYFP^ at 5 weeks of age (5 day recovery) and 12 weeks of age (7 week recovery). Tamoxifen induction for five consecutive days. The quantitative data are shown as mean ± SEM. Each dot represents data from one mouse (n=8-9). Data are from 3 individual experiments. (F) Experimental outline and frequency of EYFP^Pos^ cells at E16.5 tamoxifen labelled Cx3cr1^CreERT2^R26^YFP^ mouse pituitary gland at 7 and 14 weeks of age. The quantitative data are shown as mean ± SEM. Each dot represents data from one mouse (n=3). Data are from single experiment. (G) The cell amount of pitMØ, Ly6C^Low^MO and Ly6C^Hi^MO cells in the CCR2 KO mouse pituitary gland at 10-12 weeks of age. The quantitative data are shown as mean ± SEM. Each dot represents data from one mouse (n=5-7). Data are from 3 individual experiments. (H) Serum IL6 level in LPS and PBS (i.p.) treated mice after 24 hour recovery. The quantitative data are shown as mean ± SEM. Each dot represents data from one mouse (n=4-5). Data are from 2 individual experiments. (I) The cell amount of pitMØ, Ly6C^Low^MO and Ly6C^Hi^MO cells in the Rb^+/-^;Cx3cr1^CreERT2^; R26^EGFP^ mice mouse pituitary gland at 9 months of age. The quantitative data are shown as mean ± SEM. Each dot represents data from one mouse (n=4-10). Data are from 3 individual experiments. (J) Histological section staining of 10-month-old Rb^+/-^ mouse. Inserts are higher magnifications from the boxed anterior lobe area. Orange circle indicates excess growth of the adenohypophysis and arrows indicate abnormal vasculature. For overview images, scale is 500 µm, and for inserts, 10 µm.

**Supplemental Figure 6.**
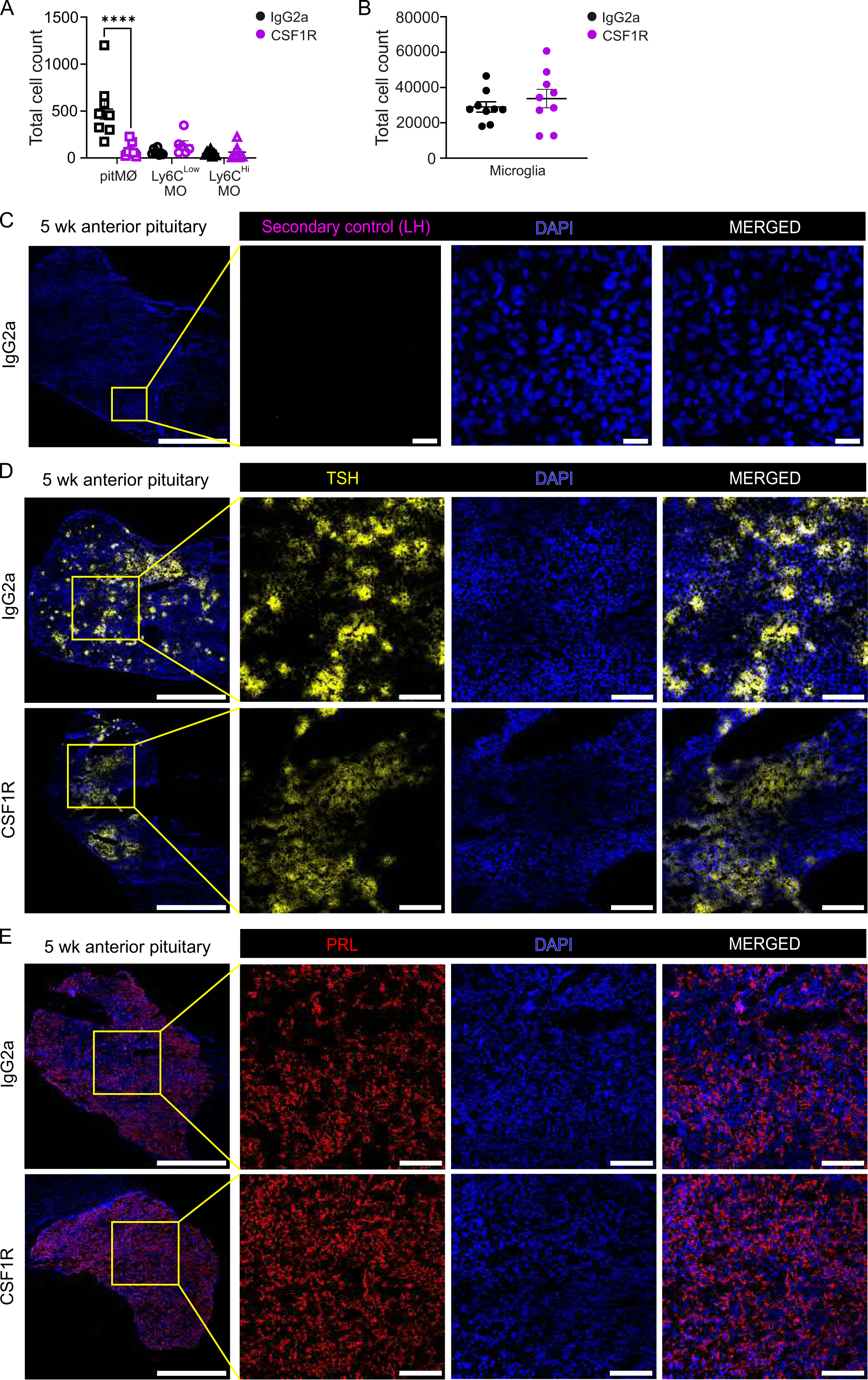
TSH and PRL in macrophage-depleted pituitary glands, related to figure 6. (A) The cell amount of pitMØ, Ly6C^Low^MO, and Ly6C^Hi^MO cells in pituitary gland of 5-week-old WT mice after treated with blocking CSF1R antibody or control IgG at E6.5, P2, P14 and 4 weeks of age. The quantitative data are shown as mean ± SEM. Each dot represents data from one mouse (n=9). Data are from 4 individual experiments. (B) The cell amount of microglia in brain of 5-week-old WT mice after treated with blocking CSF1R antibody or control IgG at E6.5, P2, P14 and 4 weeks of age. The quantitative data are shown as mean ± SEM. Each dot represents data from one mouse (n=9). Data are from 4 individual experiments. (C) Representative pituitary gland frozen section immunofluorescence staining with secondary antibody, anti-rabbit IgG goat A488, and nuclear marker DAPI from pituitary gland of 5-week-old WT mice. Inserts are higher magnifications from the boxed anterior lobe areas. For overview image, scale is 500 µm, and for inserts, 20 µm. (D, E) Representative pituitary gland frozen section immunofluorescence staining with anti-TSH (D) or anti-PRL (E) and nuclear marker DAPI from pituitary gland of 5-week-old WT mice after treated with blocking CSF1R antibody or control IgG at E6.5, P2, P14 and 4 weeks of age. Inserts are higher magnifications from the boxed anterior lobe areas. For overview image, scale is 500 µm, and for inserts, 100 µm.

**Supplemental Figure 7.**
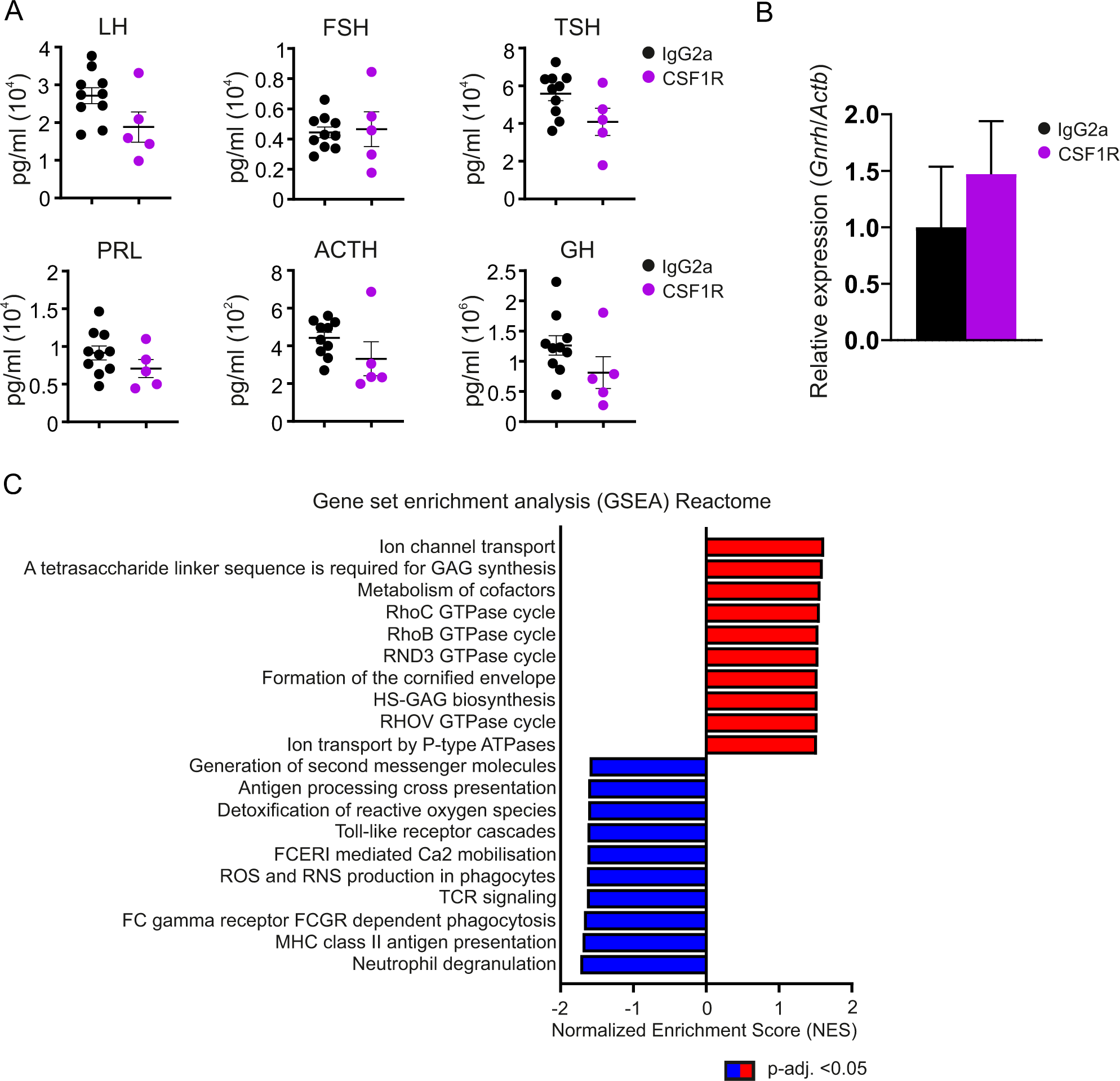
Early macrophages play a crucial role in maintaining hormonal equilibrium within the pituitary gland, related to figure 6. (A) Intra-pituitary hormone measurements for LH, FSH, TSH, PRL, ACTH and GH from pituitary gland of 4-week-old WT mice after treated with blocking CSF1R antibody or control IgG at E6.5. The quantitative data are shown as mean ± SEM. Each dot represents data from one mouse (n=5-10). (B) Relative *Gnrhr* expression in pituitary gland of 5-week-old WT mice after treated with blocking CSF1R antibody or control IgG at E6.5, P2, P14 and 4 weeks of age. The quantitative data are shown as mean ± SEM. Each dot represents data from one mouse (n=4). (C) Gene set enrichment analysis (GSEA; Reactome) of the bulk RNA-seq data. Gene sets are ranked according to their normalized enrichment scores (NES).

Table S1. Excel file containing DEGs in macrophage and monocyte clusters, Related to Figures 1 and S1

Table S2. Excel file containing DEGs in the pituitary glands of CSF1R antibody vs. control IgG treated mice, related to figures 6 and S7

Table S3. Excel file containing Ingenuity Canonical Pathways analyses of DEGs in the pituitary glands of CSF1R antibody vs. control IgG treated mice, related to figures 6 and S7

